# Teneurin-4 switches between self-recognition and canonical Latrophilin binding to direct neuronal migration

**DOI:** 10.1101/2025.09.09.671438

**Authors:** Miguel Berbeira-Santana, Claudia Peregrina, Kosuke Okuda, Jin Chuan Zhou, Maria Carrasquero-Ordaz, Amy V. Roberts, Anne E. Thomas, Evert Haanappel, Matthieu Chavent, Kamel el Omari, Lindsay A. Baker, Daniel T Pederick, E. Pardon, J Steyaert, U. Valentin Nägerl, Daniel del Toro, Elena Seiradake

## Abstract

Cortical migration is a complex process in which neurons migrate along radial glial cells (RGC) to form functional layers. Teneurins (Ten1-4) play a role by interacting with Latrophilins (Lphn/ADGRL1-3). Teneurins are also known as cell adhesion molecules, but how homophilic and heterophilic Teneurin interactions are integrated is unknown. Here, single-particle-cryo-EM data of Ten2 shows that canonical Latrophilin-binding is sterically incompatible with Ten2-dimerisation, making these interactions exclusive. We engineered surface mutations that specifically disrupt Ten2-Ten2 or Ten2-Latrophilin interactions. These are transferrable to Ten4, suggesting conserved binding mechanisms. Proteomics*, in-vivo*-gene-editing and super-resolution-microscopy show that Ten4 is expressed along RGC fibres and that migrating neurons switch from low-to-high Ten4-expression. Ten4 expression is highest in the cortical plate where Ten4-Ten4 interactions reduce RGC-attachment. In the intermediate zone, Ten4-Latrophilin interactions are required to promote neuron-RGC association. The results show how Ten4 orchestrates cortical migration by exclusive structural mechanisms, underpinning the integration of distinct migration programmes.

*Note: the adhesion GPCR ADGRL is largely referred to as ‘Latrophilin’, which is in line with previous papers in the Teneurin field. We would be happy to implement a different naming scheme if recommended*.

## Introduction

During cortical development, young pyramidal neurons migrate radially from the germinal layer in the (sub)ventricular zone (SVZ/VZ) and through the intermediate zone (IZ) to reach the cortical plate (CP), where they form cortical layers. During the initial phase of migration in the intermediate zone, neurons are multipolar with short processes, characterized by random and low speed movements along the radial axis^1^. These neurons transition to a bipolar morphology in the upper portion of the intermediate zone, close to the cortical plate boundary, where they interact with radial-glial-cell (RGC) fibres and switch to faster, fibre-guided migration before settling into functional layers in the cortical plate^2^. A cell surface receptor interaction network involving Teneurins has emerged as a key regulatory system of this migration^3^. Teneurins are large (∼2,800 amino acids) type II membrane receptors that evolved by a horizontal gene transfer event in which a bacterial toxin-like protein gene fused to the C-terminus of a eukaryotic membrane receptor^4^. Modern Teneurins are found in certain species of single-celled choanoflagellates and in bilaterians^4–8^. Mammals contain four isoforms (Ten1-4), all composed of an N-terminal intracellular domain, a single transmembrane helix, and a large ∼2,400 amino acid extracellular region. This large extracellular region is composed of a ∼200 amino acid juxta-membrane domain followed by eight epidermal-growth-factor (EGF) domains, a cysteine rich region, a transthyretin-like (TTR) domain, and a characteristic Teneurin ‘superfold’ at the C-terminus (Fig 1A). The superfold contains a spiralling beta-barrel tyrosine-aspartate repeat domain (YD-shell), which is decorated by four smaller domains: a fibronectin-like domain (FN-plug) and an ‘NCL-1, HT2A, and Lin-41’ (NHL) domain at the N-terminal end, and an antibiotic-binding domain fold (ABD) plus C-terminal DNAse-like domain (Tox-GHH)^7–11^ at the C-terminus. The amino acid chain folds into a knot-like structure: the C-terminal end of the YD-shell folds deep into the YD-shell lumen and then leads through its wall to form the ABD/Tox-GHH domains external to the YD-shell. Teneurins present two well-characterised alternative splicing sites located between the exons encoding for the EGF7 and EGF8 domains (splice site A) and in the NHL domain (splice site B). These splice sites regulate the homophilic (Ten-Ten) binding of Teneurins in cell-cell adhesion, with an insert required at either position A or B to promote cell-cell adhesion *in vitro*^9,12,13^. Teneurins also form disulphide bonded homodimers, using lone cysteines in the second and fifth EGF domains^14^. Their function as synaptic cell adhesion molecules was first described for fly Teneurins in olfactory synaptic partner matching^15–17^. A role for homophilic Ten3 interaction in murine hippocampal wiring is controversial ^12,18,19^. While there is no definitive structural model of how Teneurins interact homophilically *in trans* (from one cell to another), structure-based hypotheses have been put forward for vertebrate Ten3, Ten4 and fly Ten-m^9–11^. These published Teneurin dimer structures are different from each other, and it is unclear if any represent ‘trans’ interactions in cell-cell contacts.

**Fig 1.**
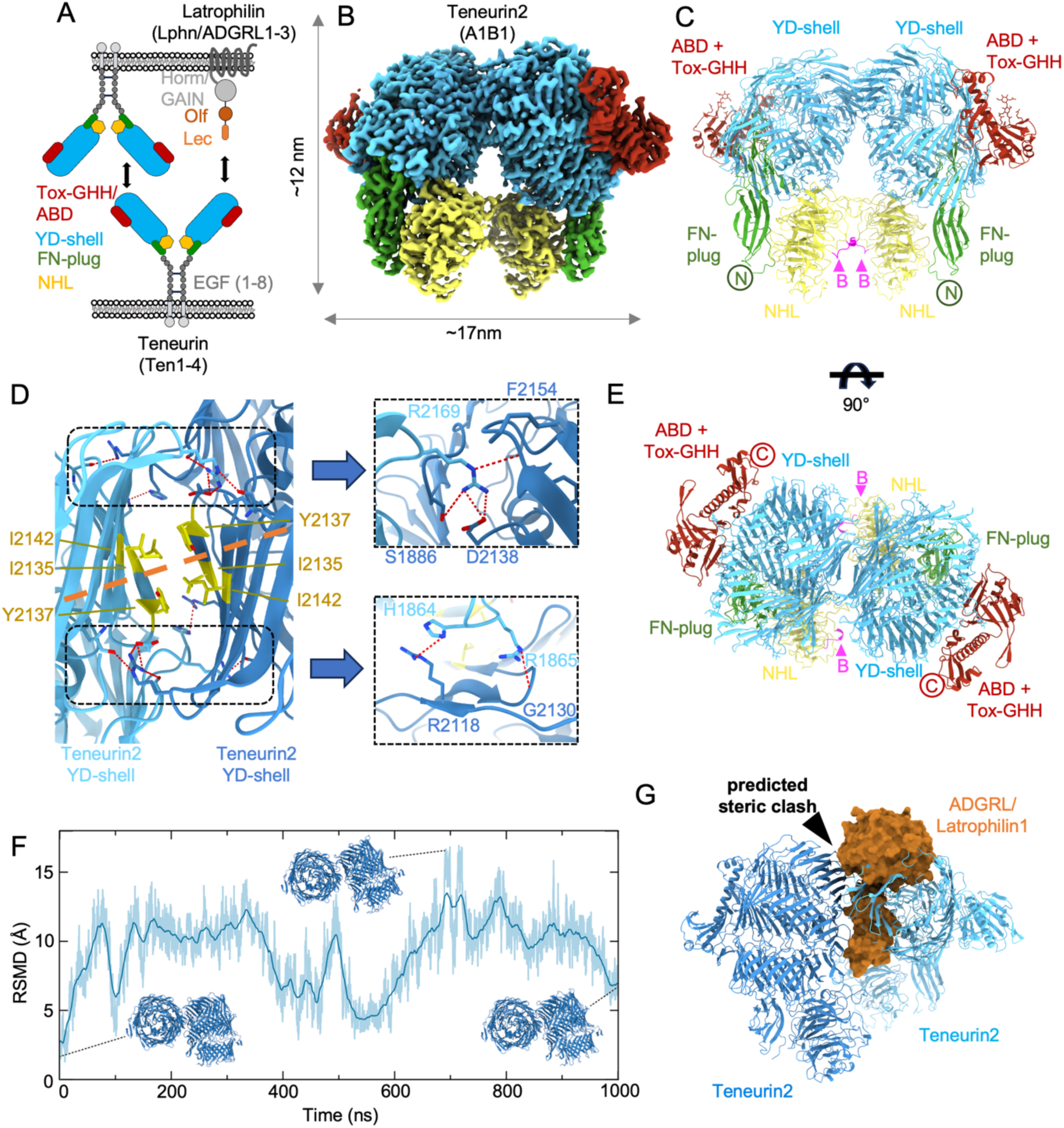
The murine Ten2 YD-shell forms a compact dimer that excludes Latrophilin-binding. **A**: A cartoon overview showing that Teneurins engage in homophilic and heterophilic interactions. Domains of the Teneurin ‘superfold’ and the Latrophilin N-terminal domains Lec and Olf are coloured separately. **B:** CryoEM density map calculated for the ectodomain of murine Ten2 isoform A1B1 after refinement. Domains coloured as in panel A. **C:** Cartoon view of the Ten2 (A1B1) dimeric model, domains annotated and coloured as in panels A and B. N termini are indicated. The location of the alternatively spliced site B (RNKDFRH in Ten2) is indicated and coloured magenta. **D:** Left: zoomed view of the Ten2-Ten2 interface showing selected residue conformations after molecular dynamics (MD) simulation with backbone constraints. Key hydrophobic residues are shown as sticks, coloured in yellow. Residues providing stable hydrogen bonds or salt bridges are shown as blue sticks. Selected atoms are coloured by type: oxygen – red, nitrogen – blue. Hydrogen bonds are indicated by thin red dashed lines. The 2-fold pseudosymmetry axis is indicated as a thick orange dashed line. Right: close views on selected areas of the interface. **E:** As panel C, but rotated by 90 degrees around the x axis. Visible C-termini are indicated. **F:** We calculated the relative root mean squared deviations (RMSD) between the YD-shell backbones of the two Ten2 copies within the dimer of the A1B1 isoform, over the course of a 1000-nanosecond (ns) unconstrained MD simulation. The dark blue line shows average values across 10 sliding window frames. The light blue coloured area depicts the spread of the RMSD values of these frames for each time point. Snapshots of the model are shown for three time points. **G:** Structural overlay of the Ten2 dimer (shades of blue) and the previously published structure of chicken Ten2 in complex with Lphn1 (orange, PDB 6SKA). The two interactions are sterically exclusive.

Teneurins also engage in heterophilic interactions such as with the adhesion G-protein-coupled receptor (aGPCR) family of Latrophilins (Lphn/ADGRL1-3, Fig 1A)^20^, the only other known binding partner in cortical migration. Latrophilins function broadly in the nervous systems of vertebrates and invertebrates, with emerging roles in mechanosensation^21–25^, and neural wiring^18,20,23,26–31^. The Teneurin-Latrophilin complex further interacts with Fibronectin Leucine-Rich Transmembrane proteins (FLRTs)^3,32–35^ via Latrophilin. The Latrophilin extracellular region contains an N-terminal lectin (Lec) domain, followed by a linker, an olfactomedin (Olf) domain and hormone (Horm) plus GPCR-autoproteolysis-inducing (GAIN) domains. We and others recently showed that the Lec domain binds to the Teneurin YD-shell and that mutations in this interface reduce Teneurin-Latrophilin binding^3,36^. Latrophilins contain a well-characterised alternatively spliced site (SS) in the linker between the Lec and Olf domains, whose presence (SS+) or absence (SS-) might modulate Latrophilin functions ^29,36^. Both Teneurins and Latrophilins are implicated in a range of neurodevelopmental diseases and other disorders such as attention deficit hyperactivity disorder, bipolar disorder and schizophrenia ^37–41^.

The wealth of recent data, while detailing different protein interaction interfaces, has failed to address how Teneurins coordinate their dual functions as both homophilic binders and receptors for Latrophilins. The lack of molecular tools to unravel these functions in an *in-vivo* context where both are simultaneously expressed, has made previous analyses challenging, reflecting a general lack of information on how multifunctional receptors integrate their functions at the molecular-to-tissue levels. Here, we use single particle cryo-electron microscopy (cryo-EM) to demonstrate that the extracellular domain of murine Ten2 dimerises in an arrangement that is distinct from previously solved structures for other Teneurins. Structural comparison with previous results ^3,36^ suggests that this arrangement clashes with Latrophilin Lec-binding at the canonical Teneurin-Latrophilin binding site. These structural insights, supported by molecular dynamics (MD) simulations, suggest that Teneurin homophilic and heterophilic interactions are structurally exclusive. We use these results to engineer Ten2 and Ten4 mutants that selectively inhibit either Teneurin-Teneurin or Teneurin-Latrophilin interactions *in trans*, and characterised a novel nanobody that selectively targets Ten4. Armed with these tools, we discovered that Ten4 is preferentially expressed along RGC fibres and that migrating neurons upregulate Ten4 as they migrate from the intermediate zone to the cortical plate. We find that Ten4-Latrophilin interactions are essential for association of neurons with RGCs in the intermediate zone, while high Ten4-expressing cells engage in homophilic, cell-repulsive interactions in the cortical plate, facilitating rapid migration along RGC fibres. The study illustrates how Ten4 uses the same interface to control both homophilic and heterophilic interactions sequentially to determine neuron-RGC attachment at different stages of cortical migration.

## Results

### Ten2 YD-shell homophilic binding clashes with Latrophilin Lec-binding

We produced full extracellular domains of the four major murine isoforms of Ten2 (A0B0, A1B0, A0B1, A1B1) using recombinant expression in mammalian cells, and performed single particle cryo-EM to determine the structures of Ten2 homodimers (Extended Data Fig. 1A). Maps after refinement have overall resolutions of ∼2.5Å (A0B0, A1B1, A1B0) –3.5Å (A0B1) and are strikingly similar for the four isoforms (Extended Data Fig. 1B-E). Key differences are found in the area that corresponds to the NHL domain, where the B1 splice form is located. In this area, the maps of those isoforms that include the B1 splice insert (A0B1, A1B1) have the highest definition, while the A0B0 map is least defined. We performed further 3D refinement for the A1B1 and A0B0 isoforms to improve density for the NHL domain, using ∼15% of the particles from each dataset (Fig. 1B and Extended Data Fig. 1F,G). We found that the map belonging to the A1B1 dataset has the best definition in the NHL region (Extended Data Fig. 1H), and so the models shown in most figures correspond to the A1B1 isoform of Ten4, unless indicated otherwise. Structural analysis resulted in a model that encompasses all of the superfold domains (Fig. 1C-E). Upstream domains, such as the EGF repeats and juxtamembrane regions, while present in the protein samples, are not resolved in any of the isoform datasets and may be flexible or flexibly linked. A structural alignment of the models derived for the four isoforms results in a root mean square deviation for C-alpha atoms (RMSD_Cα_) = ∼0.3 Å, for residues C1019-E2770 (A1B1 notation, excluding NHL domain). For all four isoforms, the YD-shells belonging to two different chains come together at a ∼75° angle, in a complex that is unlike other Ten dimer structures (Extended Data Fig. 1J,K). The surface area buried at this position in the interface is ∼2100 Å^2^, and the high resolution of the map allows for confident placing of the side chains in this interface (Extended Data Fig. 1I). We performed an all-atom molecular dynamics simulation for one microsecond to assess the stability of individual atomic interactions within the Ten2-Ten2 binding interface, using the A1B1 isoform model. Based on our established approach^3,42,54^, we imposed position restraints on backbone atoms (C, C_α_ and N), but not on side chain atoms. Taken together, the structure and atomistic simulation result suggest that the Ten2-Ten2 binding site is pseudosymmetric and consists of a hydrophobic core, which is flanked by four regions forming hydrogen bonds and salt bridges across the two monomers (Fig. 1D and Extended Data Fig. 2A-D). Two of these regions are dominated by R2169 coming from each of the monomers, and interacting with the nearby backbone oxygen of F2154, the side chain of D2138 and the backbone oxygen of S1886. The other two regions involve R1865 and R2118 from each monomer, interacting with the backbone oxygens of G2130 and side chains of H1864, respectively (Fig. 1D and Extended Data Fig. 2A).

To further assess the stability of the Ten2-Ten2 dimer, we performed an additional non-constrained simulation in which both monomers were allowed to move freely. We measured the RMSD between the backbones of the two YD-shells in the dimer, and although the dimer arrangement is stable, the RMSD values fluctuate slightly across the simulation, suggesting that the structure can “breathe” (Fig 1F). Among the most stable hydrogen bond interactions are those mediated by Ten2 R2169, while other interactions (around R1865 and R2118) are transiently broken (Fig. 1D and Extended Data Fig. 2C,D). The central hydrophobic binding surface remains relatively stable during the simulation (Extended Data Fig. 2b). In summary, the interface may open up on one side, like a hinge, conceivably allowing other binding partners to interfere and displace a Teneurin monomer.

Previously solved Ten2-Latrophilin complex crystal structures had revealed a binding site for the Lec domain of Latrophilin on the YD-shell of Ten2^3^. Structural comparison shows that this binding site, and the homophilic interaction surface we discovered here, are partially overlapping. This overlap results in a clash when the two structures are overlayed (Fig. 1G). Both binding sites localise to a similar surface region on the YD-shell including also the R1865/R2118 binding area (Fig. 2A and Extended Data Fig. 2A-D). Therefore, the two interactions are likely structurally exclusive such that Latrophilin binding could compete with Ten2 homophilic dimerisation. The dimer structures of Ten3 and Ten4 previously solved by cryo-EM^9,10^ do not engage this binding surface (Extended Data Fig. 1J,K).

**Fig 2:**
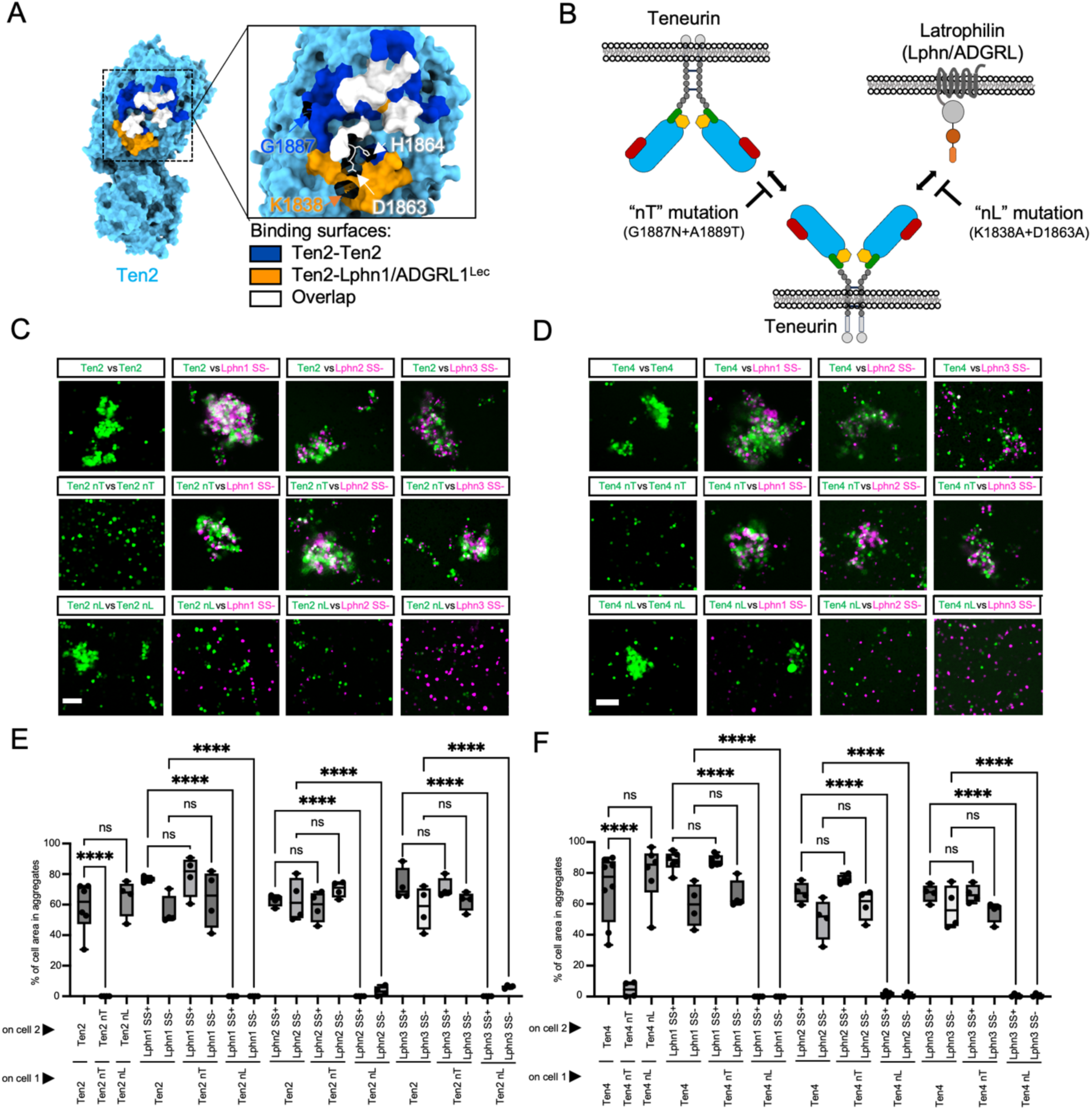
Design of Ten2 and Ten4 mutants with impaired *in-trans* binding capabilities for homophilic interaction (nT) and Lphn-binding (nL). **A**: Surface view of one Ten2 monomer. The Ten2-Ten2 and Ten2-Lphn binding surfaces are coloured separately (dark blue and orange, respectively). Residues found in both binding sites are coloured in white. Interface assignment done with PISA^99^. **B:** Summary of the properties of mutants nL and nT. **C-D:** Representative images of selected cell aggregation experiments using Ten2 **(C)** and Ten4 mutants **(D)**. Teneurin-expressing cells are shown in green, Latrophilin-expressing cells are shown in magenta. Two isoforms of Latrophilin were used (SS+ and SS-). **E-F:** Quantification of cell aggregation experiments as those shown in panels C, D. N>=4 replicates, 6 pictures per replicate. n.s. = not significant. ****p < 0.0001. One-way ANOVA test with Tukey’s post hoc analysis (E,F). Scale bars represent 100 μm (C,D).

### Structure-based protein engineering specifically disrupts homophilic or heterophilic interactions of Ten2 and Ten4

Using the structural data and molecular dynamics simulations as a guide, we designed mutants aimed at specifically disrupting Teneurin-Teneurin or Teneurin-Latrophilin binding and assessed these using cell aggregation assays. G1887N + A1889T mutations, which introduce an N-linked glycosylation site at G1887 (Fig. 2B and Extended Data Fig. 2E-H), specifically disrupt aggregation between Ten2-expressing cells, without affecting Latrophilin binding (Fig. 2C,E and Extended Data Fig. 2I-K).

Conversely, the mutations K1838A+D1863A disrupted the interaction with Latrophilins specifically, keeping the homophilic binding intact (Fig. 2C,E and Extended Data Fig. 2I-K). These mutants are termed ‘nT’ and ‘nL’, as they are ‘non-Teneurin binding’ and ‘non-Latrophilin binding’, respectively. The equivalent mutations R1881N+N1883T (nT) and R1832D+D1857A (nL) produced similar results for Ten4, showing that the interfaces are conserved in different Teneurin isoforms (Fig. 2B-F and Extended Data Fig. 2I-K). In agreement with previous studies, overexpression of wild type Ten2 or Ten4 isoforms A1B0, A0B1 and A1B1 lead to robust cell-cell aggregation, which indicates effective *trans* interactions ^3,9,12^ (Extended Data Fig. 2I,J), but we detected no mixed aggregates with cells expressing different Teneurin homologues (Extended Data Fig. 2L-N), suggesting that Teneurin homophilic interactions are homologue-specific. We also tested representative Ten3 isoforms (A0B0 and A1B1), but not Ten1 constructs, as these do not express at high enough levels at the cell surface of our *in vitro* systems, noting that Ten1 is also not very highly expressed during cortical migration (see mRNA expression data analysis below). The previously published chicken Ten2 “LT” mutation^3^, which is located in the shared region of the Latrophilin– and homophilic binding surfaces (H1864N+K1866T in Ten2 A1B1, H1858N+K1860T in Ten4 A1B1), abolishes both Teneurin-Teneurin and Teneurin-Latrophilin-dependent cell aggregation (Extended Data Fig. 3A-C), further confirming the overlapping of the two binding sites. Cell surface expression levels are similar for our mutants and wild type proteins (Extended Data Fig. 3D-F).

We also used cell-based binding assays in which purified soluble Latrophilin1-3 Lec+Olf domains interact with cell surface Teneurins. These experiments confirm that Teneurin wild type and nT mutants bind Latrophilins, and that the nL mutants do not (Extended Data Fig. 3G-J). *Vice versa*, the wild type or nT mutants of purified Ten2 and Ten4 extracellular domains bound to cell surface Latrophilins, but not the Ten nL mutants (Extended Data Fig. 3K,L). The results confirm our conclusion that wild type and nT proteins interact with Latrophilin, but not the nL mutants. Note that Teneurin-Teneurin *trans* interactions cannot be assessed by the later cell-binding method, likely because robust binding requires avidity through cell surface presentation of the receptor.

The cell surface expression of Ten2 and Ten4 constructs used in these assays was validated using immunostaining and surface biotinylation methods (Extended Data Fig. 3D-F,I,J). The specific functionalities of the nT and nL mutants are summarised in Fig. 2B.

### Characterisation of a tetramerised anti-Ten4 nanobody: NanoTen4

We produced an anti-Ten4 nanobody after immunising llamas with purified Ten4 extracellular domain protein (see methods and Extended Data Fig. 4A,B). Using a previously established approach ^42^, we biotinylated the nanobody at the C-terminus and used fluorescent Streptavidin (Atto647N/AlexaFluor^TM^633) for tetramerization (Fig. 3A,B). The tetramerised nanobody is referred to as NanoTen4. We assessed the specificity and measured the affinity of the nanobody using enzyme-linked immunosorbent assays (ELISA) (Fig. 3C,D) and we used a cell-based binding assay to assess its specificity in a cellular context (Fig. 3E,F). To test the efficacy and specificity of NanoTen4 in tissue samples, we applied it to E16.5 murine brain slices containing electroporated cells (GFP control or Ten4 knock down, see details below). In these experiments, the knocked-down neurons bound NanoTen4 significantly less than control neurons (Fig. 3G,H). We also double-stained wild type brain slices using an *in situ* hybridisation probe against Ten4 mRNA and found that its expression generally correlates with NanoTen4 labelling across different brain regions (Extended Data Fig. 4C,D).

**Fig. 3.**
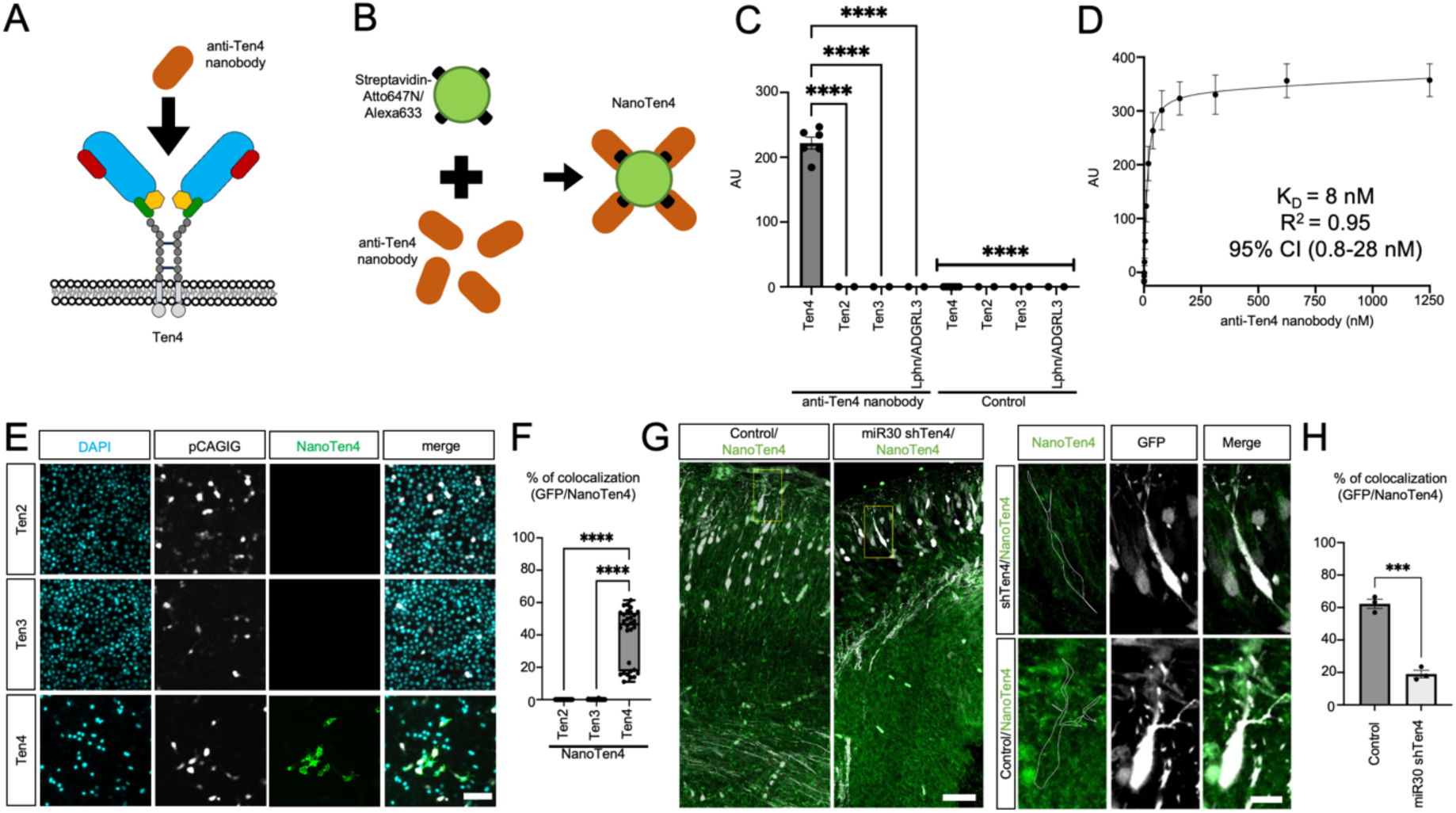
NanoTen4 specifically labels Ten4 *in vitro* and in brain tissue. **A**: An anti-Ten4 nanobody was generated to target the Ten4 extracellular region. **B:** NanoTen4 is a tetramerised form of anti-Ten4 nanobody, using fluorescently labelled streptavidin. **C:** ELISA assay shows binding of anti-Ten4 nanobody to Ten4, but not Ten2 or Ten3. N>2 independent replicates, 2 wells per replicate. **D:** Titration ELISA results suggest a Kd of ∼8 nM for binding between the nanobody and Ten4 extracellular domain. N=3 replicates, 4 wells per replicate. **E:** A cell-based binding assay confirms that NanoTen4 (AlexaFluor-633, shown in green) binds to Ten4 but not Ten2 or Ten3 on cells (GFP+, shown in white). **F:** Quantification of NanoTen4 cell-binding assays in E. N=3 experimental repeats, 10 areas per repeat. **G:** E16.5 brain slices, immunolabelled with NanoTen4 (Atto647N, shown in green). Confocal microscopy analysis shows that NanoTen4-labelling is significantly lower in GFP+ neurons, in which Ten4 expression is knocked down with siRNA (see Fig. 5 and S5). **H:** Quantification of experiments shown in panel G. n.s. = not significant. ***p < 0.001 ****p < 0.0001. One-way ANOVA test with Tukey’s post hoc analysis (C,F). Student’s t test (H). Scale bar represents 100 μm (E), 50 μm (G, left), 10 μm (G, right/inset).

### Ten4 expression is increased in the cortical plate and RGC fibres

Published single-cell RNA sequencing (sc-RNAseq) data^43^ suggests that Ten4 is the most abundantly expressed Teneurin in migrating neurons (MN) and apical progenitors (AP), including radial glial cells (RGC), at embryonic day 15 (E15, mid-gestation) (Figs. 4A and Extended Data Fig. 5A-C). To detect Ten4 protein levels in the different cortical layers (cortical plate (CP), intermediate zone (IZ), and (sub)ventricular zone (SVZ/VZ)), we dissected wildtype brains and performed tissue mass-spectrometry of the different layers separately (Fig. 4B,C). We confirmed the precision of the dissection by analysing the presence of known markers for each layer (Extended Data Fig. 5D). We found that Ten4 is the highest expressing Teneurin in the cortex, which is in agreement with the scRNA-seq analysis. Within the E15.5 cortex, we found that Teneurins are most highly expressed in the cortical plate (Fig. 4C). We also found that the three Latrophilins are present in all layers, especially the cortical plate (Extended Data Fig. 5E). For better spatial precision, we assessed Ten4 protein and RNA levels also using NanoTen4 labelling and *in situ hybridisation* (Fig. 4D-F). We found increasing Ten4 protein and RNA levels from the (sub)ventricular zone to the cortical plate, with the latter containing ∼60% of the total nanobody and RNA staining. Interestingly, confocal Airyscan imaging showed that NanoTen4 staining overlaps with Brain Lipid Binding Protein (BLBP) staining, suggesting Ten4 expression along RGC fibres (Fig. 4H). To achieve higher resolution information on Ten4 protein localisation within the intermediate zone and cortical plate, where most Ten4 is expressed, we performed Stimulated Emission Depletion (STED) super-resolution microscopy on brain samples stained with NanoTen4 and anti-BLBP. We found that Ten4 is indeed enriched close to RGC fibres (within 1.5 μm) compared to surrounding regions (Fig. 4I-K and Extended Data Fig. 5F-H), both in the intermediate zone and in the cortical plate. However, compared to the intermediate zone, more Ten4 is also expressed in non-RGC proximal areas of the cortical plate (Fig. 4K). This result agrees with the proteomics and other imaging results, suggesting that more Ten4 is expressed by neurons in the cortical plate (Fig. 4K).

**Fig. 4.**
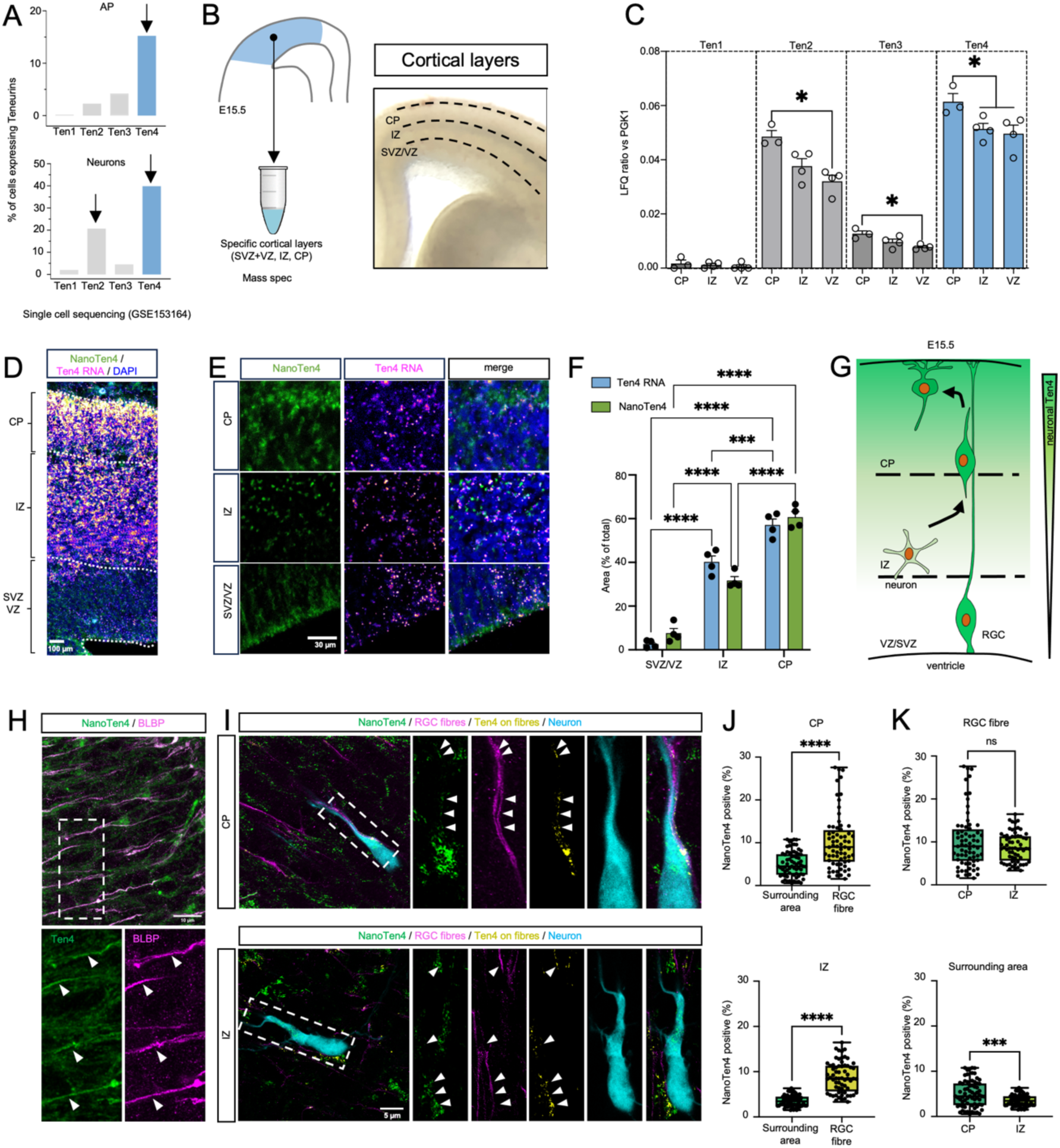
Ten4 is expressed in cortical neurons and progenitor cells, concentrated at RGC fibres, and upregulated in cortical plate neurons. **A**: Percentage of cells expressing Ten1-4 mRNA in neurons and apical progenitors (AP) based on scRNA-seq data (GSE153164)^43^. **B:** Diagram of the proteomics approach used to identify Teneurin expression levels in different cortical layers. CP: cortical plate, IZ: intermediate zone, (S)VZ: (sub)ventricular zone. **C:** Mass spectrometry results show Ten1-4 protein levels in the different cortical layers indicated in panel B. N=4 samples/group, 3 brains per sample. D, E: *In situ* hybridisation highlights Ten4 mRNA (magenta) and NanoTen4 labelling for Ten4 protein (Atto647N, green) in E15.5 cortical tissue. **F:** Quantification of data shown in panel E. N=at least 3 sections/group from 2 mice. **G:** Summary diagram indicating that Ten4 expression increases from the (sub)ventricular zone to the cortical plate. **H:** Confocal Airyscan images of E16.5 cortical plate tissue suggest spatial correlation between RGC fibres (AlexaFluor594, shown in magenta) and NanoTen4 (Atto647N, green) staining. **I:** STED images of E16.5 cortical tissue labelled with NanoTen4 (Atto647N, green) and anti-BLBP (magenta), which labels RGC fibres. A confocal image of a GFP-positive neuron (cyan) is overlayed. We used sparse GFP labelling and therefore other neurons are not visible in this image. Yellow indicates Ten4 protein present on RGC fibres. Panels are a maximum projection of two slices of a Z-stack. **J,K:** Quantification of data shown in panel I, for cortical plate (CP) and intermediate zone (IZ). Areas within a 1.5 μm radius of the BLBP-positive fibres versus the surrounding tissue were analysed separately (’RGC fibre’), and the percentage of NanoTen4-positive pixels within these area versus the ‘surrounding areas’ was quantified (panel J). Panel K is based on the same quantification results as used in panel J, but comparing values between the IZ and CP. N=1 brain slice, >64 pictures per replicate. n.s. = not significant. *p < 0.05 ****p < 0.0001. One-way ANOVA test with Tukey’s post hoc analysis (C,F). Student’s t test (J,K). Scale bars, 100 μm (D), 30 μm (E), 10 μm (H), and 5 μm (I).

Taken together, while Ten4 is preferentially expressed at RGC fibres across the cortical plate and intermediate zone, migrating neurons seem to upregulate Ten4 expression upon reaching the cortical plate.

### Cortical migration requires homophilic and heterophilic Ten4 interactions at different stages

To study the distinct roles of Ten4 interactions *in vivo* we used an established murine *in-utero* electroporation (IUE) model for genetic manipulation at embryonic age E13.5. The resulting tissues are analysed three days later at E16.5 (Fig. 5A,B). Using this method, we show that knock-down^3,42,44^ of Ten4 with two short-hairpin RNA (shRNA) delayed the migration of neurons within the cortical plate, so that fewer neurons reached the upper layer compared to the control (Fig. 5C,D and Extended Data Fig. 6A-E). An experiment where we overexpressed wild type Ten4 resulted in an even stronger delay, with most neurons remaining in the intermediate zone, failing to enter the cortical plate (Fig. 5E,F). Introduction of the mutants nT and nL in these over-expression experiments resulted in significant rescue. Of the two mutants, the nL mutant showed the most pronounced effect, with a rescue of ∼65% of cells reaching the cortical plate, compared to the less pronounced but significant rescue of ∼24% for the nT mutant (Fig. 5E,F).

**Fig. 5.**
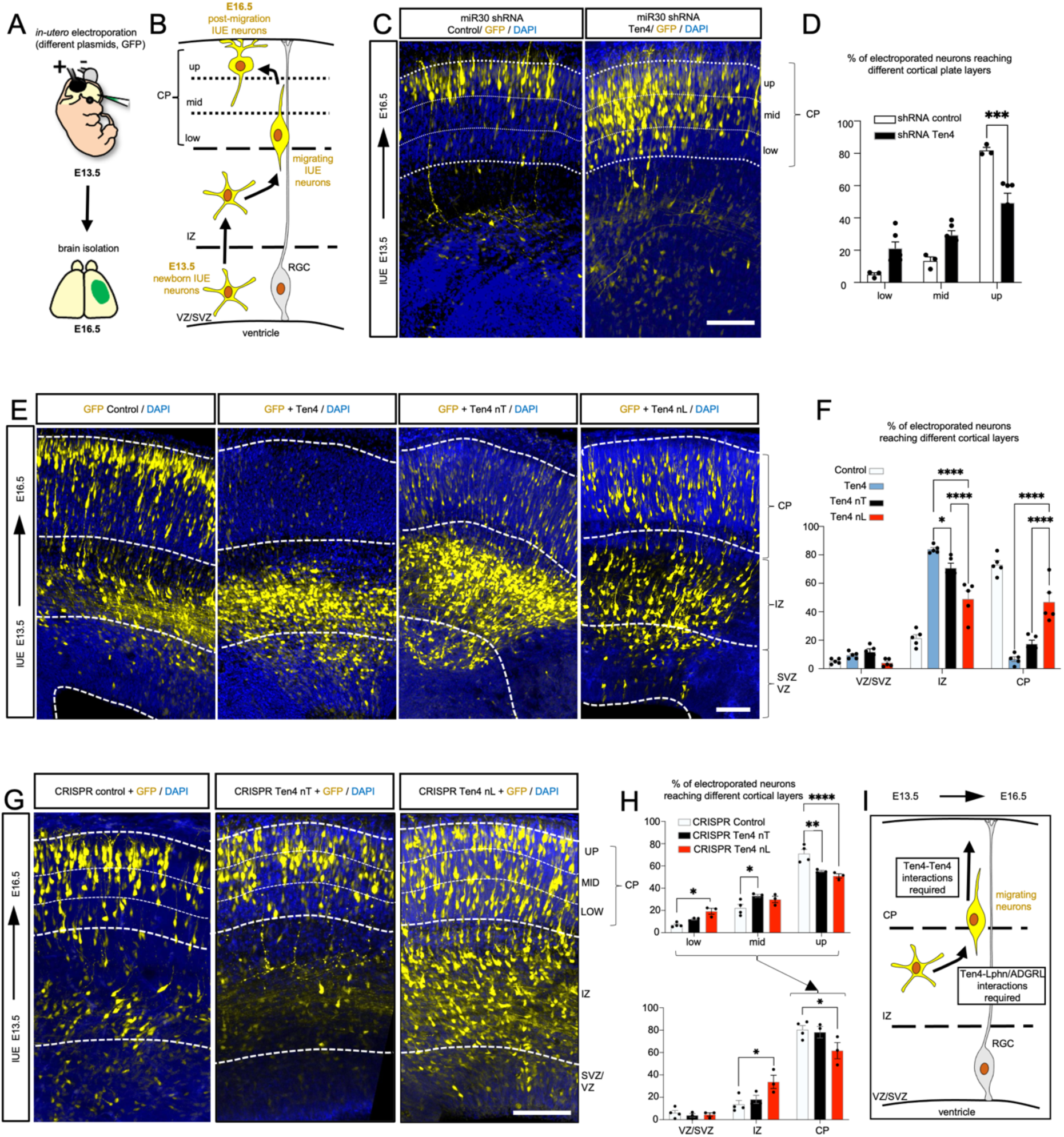
Ten4 directs different stages of cortical migration via homophilic and heterophilic interactions. **A**: Schematic of in utero electroporation (IUE) performed at E13.5. **B:** Diagram of radial migration during the course of an IUE experiment. **C:** Coronal sections of an E16.5 electroporated murine cortex. Neurons were electroporated with pCACIG and pCAG-miR30 containing two shRNAs for murine Ten4. The cortical plate was subdivided into 3 bins (up, mid, and low), and the number of GFP+ neurons in each bin was quantified. **D:** Quantification of experiments shown in panel C. N=3 electroporated brains (control), N=6 (shRNA Ten4). **E:** Coronal sections of an E16.5 cortex previously electroporated with pCAGIG (GFP) control plasmid or pCAG-Ten4-IRES-GFP, pCAG-Ten4nT-IRES-GFP or pCAG-Ten4nL-IRES-GFP expressing different Ten4 constructs in addition to GFP. **F:** Quantification of the data shown in panel E. N=5 electroporated brains for all conditions. **G:** Coronal sections of an E16.5 electroporated cortex. Neurons expressing GFP were modified with CRISPR/Cas9 to block Ten4-Ten4 interaction (Ten4 nT), Ten-Latrophilin interaction (Ten4 nL) or not modified in the *Tenm4* locus (CRISPR control). **H:** Quantification of the data shown in panel H. N>3 electroporated brains. *p < 0.05, **p < 0.01, ***p < 0.001. We performed one-way ANOVA test with Bonferronís (D, H – top panel) or Tukey’s (F, H – bottom panel) post hoc analysis. Scale bars represent 100 μm (C,E,G).

Overexpression and knock-down experiments have certain limitations: a complete knock-out/knock-down of a multifunctional receptor abolishes all its functions, thereby making it difficult to unravel the specific roles of individual interactions. Also, the phenotype associated with over-expression experiments, even with structure-based mutants as controls, are often not physiological and only valid within a rescue framework. Therefore, we used a CRISPR/Cas9-based approach based on previously published methodology^3,45^ to introduce our structure-based mutations (nT and nL) in the endogenous *Tenm4* locus. In agreement with the above described results, we find that disrupting either Ten4 homophilic or heterophilic interactions affects neuronal migration (Fig. 5G,H), but as seen before, there were differences between the two mutants: the Ten4 nT mutant cells successfully entered the cortical plate, but a significant number failed to reach the upper layer, indicating a delay in migration through the cortical plate. In contrast, the Ten4 nL mutant cells accumulated in the intermediate zone, at the boundary with the cortical plate (Fig. 5G,H). To validate that the CRISPR-induced nT and nL mutations are present in our brain slices, we used BaseScope^TM^ to directly visualize edited mRNA in GFP^+^ neurons. We found that ∼50% of the GFP^+^ neurons expressed the mutated transcript (Extended Data Fig. 6F-H). We further confirmed these results by amplicon sequencing to identify the genomic edits in electroporated tissue (Extended Data Fig. 6I,J).

In summary, these results suggest that Teneurin homo– and heterophilic interactions act at different points to direct migration through the intermediate zone and cortical plate. First, the Ten4-Latrophilin interaction promotes neuronal entry into the cortical plate. Then, higher Ten4 expression and functional Ten4-Ten4 interactions are required for effective migration through the cortical plate (Fig. 5I). Functional Ten4 homophilic interactions likely depend on the increased Ten4 levels present in neurons of the cortical plate.

### Ten4 regulates neuron attachment to RGC fibre through distinct interactions

To visualise gene-edited neurons alongside non-edited wild type neurons we applied shadow imaging^46,47^ to our CRISPR/Cas9-edited samples (Fig. 6A,B and Extended Data Fig. 7A). The technique highlights the contours of all cells by staining the extracellular matrix within a tissue ^46,47^. It is a powerful method for visualising all cells in an unbiased way and is compatible with immunostaining. We labelled RGC fibres with anti-BLBP, while GFP highlights CRISPR/Cas9-targeted neurons. In comparing gene-manipulated neurons with WT controls, we found that the introduction of the Ten4 nT and nL mutations change the association patterns with RGC fibres without affecting cell size in the cortical plate (Fig. 6C-E and Extended Data Fig. 7B). Frequency distribution analysis showed that the fibre contact area of Ten4 nT neurons varies significantly more compared to the wild type control, whereas the distribution is narrower for Ten4 nL neurons. Neurons expressing Ten4 nT have increased contacts with RGC fibres compared to control neurons, while neurons expressing Ten4 nL have reduced contacts. We also analysed neurons located in the intermediate zone, although as shown in Fig. 5, mainly Ten4 nL expressing neurons remain in the intermediate zone at the point of analysis, while Ten4 nT cells have largely migrated into the cortical plate at that point. We found no difference in cell size or RGC fibre contact area when comparing control and Ten4 nL cells in the intermediate zone (Fig. 6C,F,G and Extended Data Fig. 7B).

**Fig. 6.**
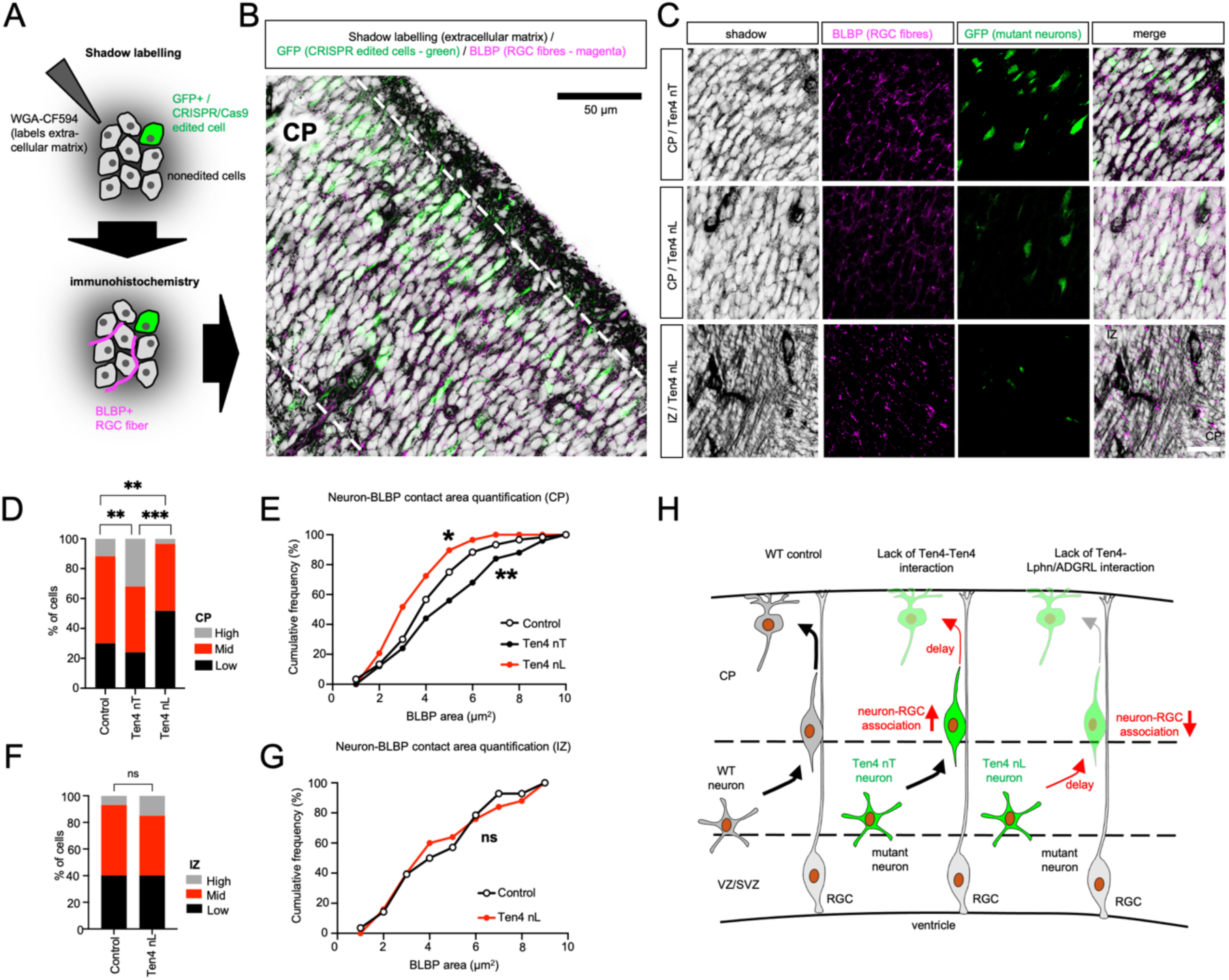
Ten4 regulates neuron-RGC association via two distinct mechanisms. **A**: Diagram depicting the shadow and immunohistochemistry labelling strategy for brain slices collected after IUE using CRISPR/Cas9 reagents to target *Tenm4* (as in Fig. 5G). Black = extracellular matrix, magenta = anti-BLBP, GFP-positive CRISPR-targeted neurons= green. Wild type neurons are white. **B:** Example of a labelled brain slice with channels merged, following the strategy outlined in panel A. **C:** Examples of single channel and merged images for each condition tested. **D-G:** Quantification of experiments as shown in panel C. We expanded the outlined somata of cells by 2 μm and quantified the BLBP labelling within these areas for different cells (control, nL and nT). The neurons were separated into different populations depending on the levels of RGC staining in their vicinity: low (0-3.5 mm^2^), mid (3.5-6.5 mm^2^), and high (6.5-10 mm^2^). The plots display the fraction of cells and cumulative frequency distributions in the CP. N=3 brains. **H:** Summary diagram showing the observed phenotypes. n.s. = not significant. *p<0.05, **p < 0.01, ***p<0.001. Chi-square contingency analysis (D,F) and Kolmogorov-Smirnov test (E,G). Scale bars represent 50 μm (B) or 20 μm (C).

The results are consistent with previous studies indicating that migration through the intermediate zone generally lacks defined polarity and is not along RGC fibres until the neurons switch to saltatory migration as they enter the cortical plate^48–50^.

Taken together, these results suggest that different Ten4 interactions direct the association of neurons to RGC fibres in distinct, even opposing ways, and that they play a role at different time points: Latrophilin interactions play a major role initially, at the border to the cortical plate, as neurons attach to the RGC fibres, while homophilic interactions regulate attachment and fast migration further up in the cortical plate. Interfering with these distinct cellular mechanisms results in different migration phenotypes (Fig. 6H).

### Ten4 upregulation in neurons acts as a switch for migration

Our results thus far have pointed to a phenotypic switch, where intermediate zone neurons expressing lower levels of Ten4 react less to externally presented Ten4, compared to cortical plate neurons expressing higher levels of Ten4. To confirm that cortical neurons express different levels of Ten4, we analysed our published scRNA-seq^87^ data derived from dissociated cortical neurons at E15.5 (Extended Data Fig. 7C). We show that Ten4 is indeed preferentially enriched in neurons populating the cortical plate such as Bcl11b+ neurons (Fig. 7A,B and Extended Data Fig. 7D). Latrophilins are not preferentially enriched in these neurons (Fig. 7C,D). Also, we found different levels of Ten4 expression across the neuronal population (Fig. 7E), while the expression levels of Latrophilins were relatively constant (Extended Data Fig. 7E). The data analysis also shows that there is very low correlation between Ten4 expression and that of other Teneurins or Latrophilins (Extended Data Fig. 7F,G). Next, we tested whether high and low-level Ten4 expressing neurons react differently when challenged with external Ten4.

**Fig. 7.**
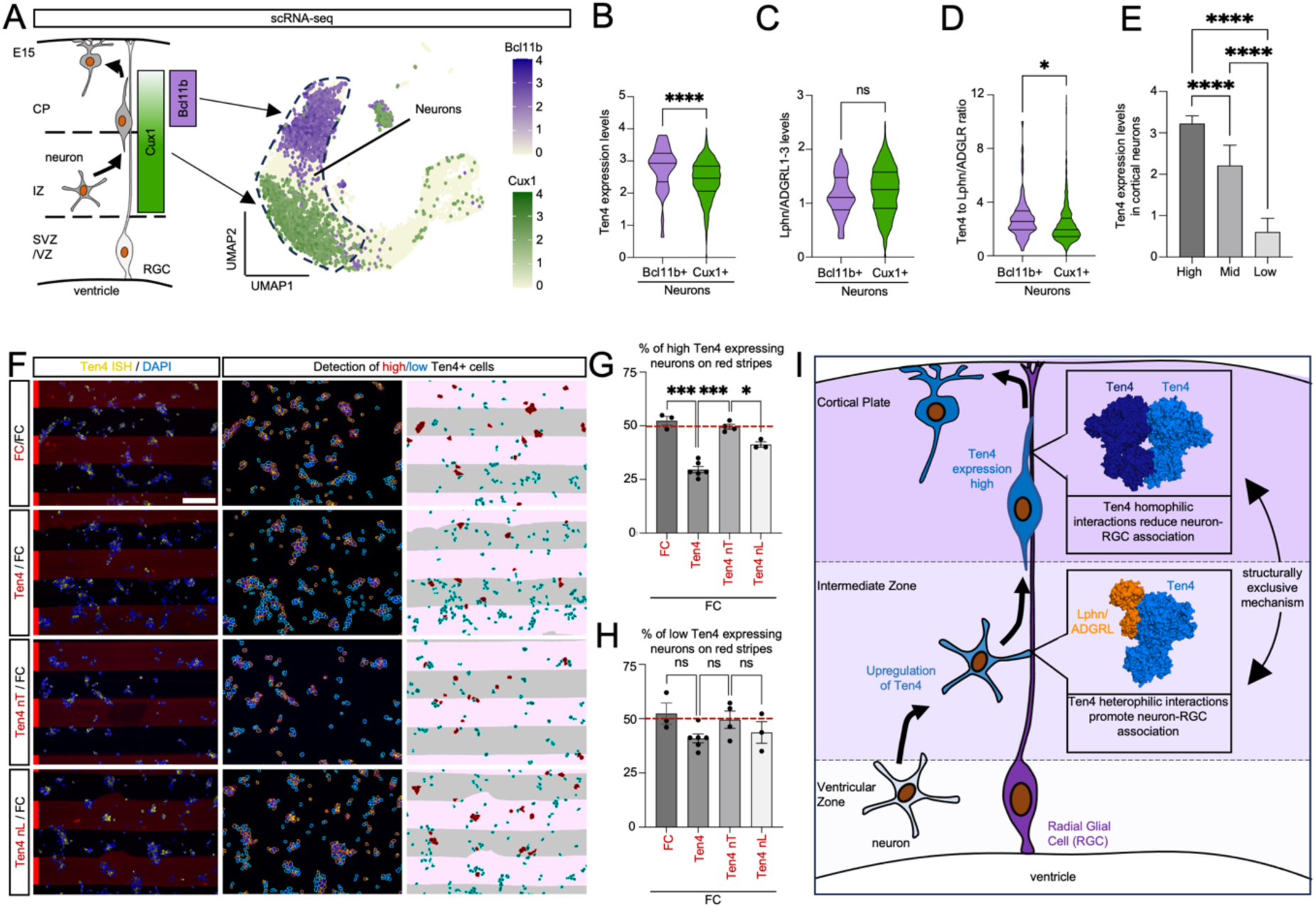
Homophilic Ten4 interaction triggers cell-repulsion in Ten4-expressing neurons. **A**: Schematic showing where the markers Bcl11b and Cux1 are expressed in the cortex, and a UMAP derived from scRNA-seq analysis of cortical cells at E15.5 (GSE271794^119^). The cells are coloured depending on their expression of Bcl11b (purple) or Cux1 (green). **B:** Levels of Ten4 expression in Bcl11b+ and Cux1+ neurons, after analysis of the data indicated in panel A. The result shows that Ten4 is predominantly expressed in neurons of the cortical plate. **C:** Levels of Latrophilin1-3 expression in Bcl11b+ and Cux1+ neurons. Latrophilins are not enriched in any particular section of migrating neurons. **D:** Ratio of Ten4 to Latrophilin1-3 in Bcl11+ and Cux1+ neurons. Ten4 is enriched in Bcl11b+ neurons compared to Lphn/ADRGLs. **E:** Cortical neurons exhibit distinct levels of Ten4 mRNA expression, here binned into high, mid and low. **F:** Representative images of a stripe assay in which E14.5 dissociated cortical neurons are challenged with WT, nT or nL Ten4 ectodomains. Neurons were grown on alternate stripes (red and black) containing Fc, Ten4, Ten4 nT or Ten4 nL. In-situ hybridisation was used to indicate Ten4 mRNA levels (yellow). Nuclear staining with DAPI is shown in blue. We used an automated detection pipeline to sort neurons into high and low Ten-expressing bins (see Fig S7I). **G,H:** Quantification of experiments shown in panel F, after binning. N=3 or more experiments per condition. **I:** Overview diagram summarising functions of Ten4 during cortical migration. n.s. = not significant, *p < 0.05, **p < 0.01, ***p < 0.001, one-way ANOVA test with Tukey’s post hoc analysis (E,G,H). Student’s t test (B,C,D). Scale bar represents 100 μm (F).

We used dissociated cortical neurons (E14.5) in a stripe assay to test whether high Ten4-expressing neurons react more to external Ten4 (Extended Data Fig. 7H,I). In these assays, migrating cells choose between surfaces containing purified Ten4 protein (WT, nT or nL) and neutral control protein (recombinant Fc fragment protein (Fc)). *In-situ* hybridization labelling of Ten4 mRNA revealed cells expressing high levels of Ten4 versus those expressing low levels (Fig. 7F and Extended Data Fig. 7I). Quantification of these experiments showed that wild type Ten4, but not the nT mutant, is repulsive for high Ten4-expressing neurons (Fig. 7F,G), suggesting that Ten4-Ten4 homophilic ‘*trans*’ interactions are repulsive in these neurons. This result is in agreement with an increase of RGC fibre attachment for neurons expressing Ten4 nT, *in vivo* (Fig. 6C-H) as these lack the ability to engage in homophilic Ten4-Ten4 interactions. Interestingly, Latrophilin interactions still play a role in these neurons as the mutant Ten4 nL protein stripes are less repulsive than the wild type protein, although still more repulsive that the nT mutant protein. In line with the model in which low-level Ten4 expressing cells do not engage in *trans* homophilic Ten4 interactions, we found that low Ten4-expressing neurons do not react significantly to Ten4 (Fig. 7F,H), suggesting that even Latrophilin signalling, if switched on, is not causing a strong effect in these neurons.

## Discussion

Brain development relies on the multiple activities of receptors and adhesion molecules that operate in a context-dependent way. Different complexes can form between receptors and their diverse ligands at different times or simultaneously in different cell-to-cell configurations. This presents a challenge to studying the functions of individual interactions *in vivo*. We have developed an integrative approach that leverages structural biology-based protein engineering and nanobody-based tools to unravel how neurons integrate different stages of radial migration using structurally exclusive Ten4 interactions (Fig. 7I). In contrast to previously determined dimer structures^9,10^, we found that Ten2 forms a distinct dimer using a binding site on the YD-shell, which overlaps with the previously determined Latrophilin-binding site (Fig. 1 and Extended Data Fig. 1). These results provide a satisfactory explanation for why some mutations in this area abolish both homophilic and Latrophilin-binding in *trans* (see^3^ and Extended Data Fig. 3A-E). They also, finally, shed light on the elusive molecular mechanisms that underpin Teneurin-Teneurin *trans* interactions.

Interestingly, the same interface was not observed in the previously published Ten4 dimer structure ^9^, even though our mutational analysis points to a conserved *trans*-binding mechanism for Ten2 and Ten4. An alignment of this previously published Ten4 dimer ^9^ and our structures is sterically compatible, forming a continuous array (Extended Data Fig. 1J), it is therefore conceivable that both are used to mediate different Ten4 functions, even simultaneously.

How cortical migration is orchestrated by different molecular interactions to allow neurons to migrate to their final targets is overall poorly understood. Here, we identify Ten4 as a key regulator during two important steps: Ten4 interacts with Latrophilin to promote RGC-attachment during the transition from the intermediate zone into the cortical plate, while during subsequent RGC-dependent migration within the cortical plate, Ten4 forms homophilic interactions to reduce RGC attachment. This functional switch is underpinned by the dynamic expression of Ten4, i.e. neurons migrating into the cortical plate upregulate Ten4 expression, and a structural switch in which Ten4-Ten4 and Ten4-Lphn binding are sterically exclusive. Our findings position Ten4 as a regulator of the attachment dynamics that neurons exert on RGC fibres as they migrate through the cortical plate. Indeed, Ten4 regulates the activity of the focal adhesion kinase (FAK)^57^, which is required for proper interaction between neurons and RGC fibres^58^. Further questions remain; could Ten4 interactions also contribute to migration termination? Interestingly, a recent study identified a repulsive interaction between Sema6A, expressed in RGC fibres, and PlxnA2/A4, enriched in neurons reaching the top of the cortical plate. This interaction acts as a termination signal by inducing permanent neuronal detachment from the RGC fibres^59^.

Moreover, increased neuron-neuron interactions are thought to weaken neuron-RGC fibre contacts during the terminal phase of migration. Given that Ten4 is upregulated in neurons within the cortical plate, it is conceivable that Ten4 could function in alternative topologies, such as mediating interactions between neurons, which are densely packed in this region. A similar mechanism has been described for integrins, high α3 integrin expression in the upper part of the cortical plate reduces neuron-RGC attachment while enhancing neuron-neuron interactions ^60^. Interestingly, previous studies suggest that Teneurins may function redundantly with integrins^61^. Upregulation of factors involved in switching migration patterns has also been observed for other proteins. As neurons migrate through the cortical plate, several axon guidance receptors are expressed, most notably members of the Eph/ephrin protein family, such as EphrinB1^51^, EphA7^52^, and EphB6^53^, which regulate the lateral dispersion and columnar distribution of neurons within this layer. Other semaphorins and their receptors, Plexins, are also enriched in the cortical plate. For example, Sema7A/4D^54^, as well as Sema3E/PlxnD1^55^, control neuron lamination.

The intricate interdependence between molecular-level and tissue-level events that we show here for Ten4, is reminiscent of Teneurin functions in hippocampal wiring. In this context, Teneurin-expressing axons first encounter high-expressing Latrophilin regions, before reaching high-Teneurin expressing areas where synapses are formed^18^. Although our stripe experiments with cortical neurons results show a cell-repulsive effect for homophilic Ten4-interactions, hippocampal axons expressing Ten3 preferentially target and synapse with Ten3-expressing dendrites, suggesting that homophilic Teneurin interactions promote cell-contacts in this context. Indeed, homophilic axonal Teneurin interactions were first described as promoting adhesion and synapse formation in different biological systems ^12,15–18^. The results illustrate that Teneurin homophilic interactions function in context-dependent ways, likely eliciting different functions in axons compared to migrating cells. This has also been shown for Teneurin-Latrophilin interactions, where, within the same experiment, Ten2-Latrophilin interaction was repulsive for cortical cells, but not for cortical axons^3^. Intriguingly, migrating cortical neurons establish transient glutamatergic synapses with subplate neurons before entering the cortical plate, and disruption of these early synapses causes cortical migration defects^67^. Therefore, Teneurins, Latrophilins, and their interaction partners, could transiently function as ‘synaptic proteins’, also during cortical migration^50^. Whether they do so, and how this integrates with other functions during migration, is a stimulating question for future research. A multifaceted role in mediating context-dependent adhesion or repulsion has been observed also for other receptors functioning in both cell migration and synapse specification, such as the Eph/ephrin protein family ^68,69^. As for Ten4, receptor concentrations can be important for Eph functions. *In vitro* assays have shown that low levels of EphA promote attraction^70^, whereas high levels induce repulsion^71^. A similar effect is observed during topographic mapping, where growth cones migrate toward the superior colliculus: low levels of ephrinA promote attraction, while high levels trigger repulsion and inhibit migration^72^. Beyond the cortex and hippocampus, Teneurins and Latrophilins are found in multiple areas of the developing brain ^27,73–77^, as well as in adult tissues and cancers^78–82^. Mutations and risk variants of Ten4 have been implicated in various brain disorders, including bipolar disorder^83^, schizophrenia^40^, mood disorders^84^, early-onset Parkinson’s Disease^85^, essential tremor^86^, disrupted axonal guidance^86^, and impaired oligodendrocyte myelination^87^, a process that is thought to depend on Ten4 homophilic interactions^88^. Dysregulated expression of Ten4 and Ten2 in particular, is implicated in several cancer types^89–98^. The mutants, nanobodies and concepts we have developed here will likely be relevant for the study of these tissues and could shed light on the associated diseases. The work introduces a robust methodological framework for understanding the role of molecular interactions withing complex tissue environments.

## Acknowledgements

We thank Dr. Rishi Matadeen, Dr. Joseph Caesar, Dr. Teige Matthew-Palmers, and Dr. Edward Lowe at the COSMIC cryo-EM facility (University of Oxford) for support with data collection and data processing. We thank María Calvo from the Advanced Microscopy service (CCiT, university of Barcelona) for help with confocal microscopy. We thank Dr. Rüdiger Klein for access to the proteomics facility of the Max Planck Institute of Biochemistry (Martinsried, Germany) and its facility members for support and sample processing. We gratefully acknowledge Dr. Lothar Schermelleh, Dr. Niloufer Irani and Dr. Deirdre Kavanagh of the Leica Microsystems Centre of Excellence at the Micron Bioimaging Facility, University of Oxford for their support and assistance in this work. M.B.-S was funded by the Pelly-Bannister Scholarship (Somerville College, Oxford). E.S. was supported by a Wellcome Trust (202827/Z/16/Z and 226647/Z/22/Z) and the EMBO Young Investigator Programme. D.d.T was funded by the Ramón y Cajal program (RYC-2017-233486), MINECO project: CNS2023-144706. C.P was funded by an FI fellowship from Generalitat de Cataluña. This work was granted access to the HPC resources of TGCC Joliot-Curie supercomputer (under the GENCI allocation A0160715135 and SS010715367). We thank Prof. Liqun Luo for providing Teneurin plasmids. We thank Dr. Marjorie Fournier and Dr. Vaishnavi Ravikumar of the Advanced Proteomics Facility of the Department of Biochemistry (University of Oxford) for the analysis and processing of the LC-MS analysis of Ten2 and Ten4 WT and TT proteins. This work benefited from access to the VIB-VUB centre for Structural Biology, an INSTRUC-ERIC centre, part of the European Strategy Forum on Research Infrastructures (ESFRI), and the Research Foundation – Flanders (FWO) for their support to the nanobody discovery. Financial support was provided by INSTRUCT-ERIC (PID: 23770). We also thank Eva Beke for technical assistance during nanobody discovery.

## Author contributions

M.B.-S. was responsible for structural biology, protein production, protein engineering, cell aggregation assays, cell-binding assays, cloning, nanobody characterisation, and STED microscopy; C.P. led the *in vivo* work, expression analyses and stripe assays; K.O. led the shadow imaging work; J.C.Z. established cryo-EM data collection and processing, and contributed to cryo-EM data processing; M.C.-O. produced vectors, protein, and contributed to nanobody production; E.H. performed MD simulations and analysis; M.C. oversaw the MD simulations and analysis; A.V.R. contributed to cloning and cell-based aggregation assays; A.E. T. contributed to cell-based aggregation assays; K.e.O. oversaw model building and refinement; L.B. contributed to cryo-EM data analysis; D.T.P. provided template Teneurin vectors; E.P. and J.S. performed nanobody discovery; V.N. oversaw the shadow imaging work; D.d.T. oversaw the *in vivo* work, expression analyses and stripe assays; E.S. oversaw structural biology, binding studies and protein engineering. All authors have contributed to the manuscript.

## Supplemental information titles and legends

Supplemental Table 1 related to Extended Data Fig. 1: cryo-EM processing statistics for the density maps.

Supplemental Table 2 related to Extended Data Fig. 1: Refinement statistics for the atomic models.

## Data Availability

The electron density maps for the mouse Ten2 dimers have been deposited in the Electron Microscopy Data Bank under the accession codes: EMD-51022 (A0B0), EMD-50975 (A1B1), EMD-50976 (A1BO), and EMD-51021 (A0B1).The atomic coordinates and models resulting from the structural analysis of the aforementioned maps have been deposited on the Protein Data Bank under the following accession codes respectively: 9G42 (A0B0), 9G2F (A1B1), 9G2H (A1B0), and 9G41 (A0B1).

## Code Availability

The scripts and macros used for quantification of the stripe assays or cell binding assays can be made available upon request to the lead contacts Daniel del Toro, and Elena Seiradake.

## Materials availability

This study generated new a new anti-Ten4 nanobody (CA20030) which can be made available upon request to the VIB (Els Pardon/Jan Steyaert) and the lead contact (Elena Seiradake). CDR loop sequences are available in the Nanobody discovery methods’ section and Extended Data Fig. 4A,B.

## Online methods

### Mouse embryos

All mice (C57BL/6 background) were housed with a 12h:12h light:dark cycle and food/water available ad libitum. All animal experiments were used in accordance with the ethical guidelines (Declaration of Helsinki and NIH, publication no. 85-23, revised 1985, European Community Guidelines, and approved by the local ethical committee (University of Barcelona, 225/17 and Generalitat de Catalunya, 404/18).

### Primary cultures

Neurons were dissociated from cortices of E14.5 embryos and cultured on stripes as described previously^3^. Neurons were cultured for 1 day in vitro at 37°C, 5% CO_2_ in Neurobasal^TM^-A medium (Invitrogen, Cat#A3582901) supplemented with B27 (GIBCO, Cat#17504044). Then cells were fixed with 4% Paraformaldehyde (SIGMA-Aldrich, Cat#158127-100G) for 10 min followed by immunostaining.

### Cell lines

K562 suspension cells (ATCC; CCL-243; RRID: CVCL_0004) were cultured in RPMI-1640 media (LGC Standards, Cat#ATCC 30-2001) supplemented with 10% FBS (GIBCO, Cat#10437028).

HEK293T cells (ATCC; CRL-3216; RRID: CVCL_0063) were cultured in DMEM plus L-glutamine (Thermo Fischer Scientific, Cat#41966052) supplemented with 10% Fetal Bovine Serum (FBS) (GIBCO, Cat#10437028), and 5% Non-Essential Amino Acids (Life Technologies, Cat#11140035). Cell lines were maintained in sterile conditions in a 37°C, 5% CO_2_-incubator.

### Vectors and cloning

We cloned constructs of mouse Ten2 (Uniprot: A0A0A0MQB7) (Ten2 A0B0, A1B1, A1B0, A0B1: residues 1-2774*; Ten2^ecto^: residues 398-2774*; nT mutant: G1887N+A1889T; nL mutant: D1863A+K1838A; LT mutant: H1864N+K1866T), mouse Ten4 (Uniprot: Q3UHK6) (Ten4 A0B0, A1B1, A1B0, A0B1: residues 1-2771*; Ten4^ecto^: residues 364-2771*; nT mutant: R1881N+N1883T; nL mutant: D1857A+R1832D; LT mutant: H1858N+K1860T), Latrophilin1 (Uniprot: H7BX15) (ADGRL1 residues 22-1471), Latrophilin2 (Uniprot: A0A0G2JGM8) (ADGRL2 residues 1-1487), and Latrophilin3 (Uniprot: Q80TS3-3) (ADGRL3 residues 1-1543). We used previously published mouse Latrophilin1-3 constructs as indicated^3^, including Lphn/ADGRL1^Lec-Olf^ (residues: 29-395, SS+, pHL-Sec C-terminal eAvi), Lphn/ADGRL2^Lec-Olf^ (residues: 30-400, SS+, pHL-Sec C-terminal eAvi) and Lphn/ADGRL3^Lec-Olf^ residues: 82-466, SS+, pHL-Sec C-terminal eAvi). For protein expression, the Teneurin ectodomain constructs were cloned into the Age1-Kpn1 site of the pHL-Sec vector (Addgene; Cat#99845)^100^. For structural studies and biotinylated nanobody production, we use a pHL-Sec vector that contains an N-terminal secretion signal peptide which is used instead of the native signal peptide of the proteins. The vector also contains a 6xHis tag and an Avi-tag (protein sequence: GLNDIEAQKIEWHE). For nanobody production in bacteria, we use a pMESy4 vector (GenBank KF415192) that contains a His-tag and EPEA-tag at the C-terminus. The nanobody was cloned behind a pelB pre-signal (MKYLLPTAAAGLLLLAAQPAMA) that directs the nanobodies to the periplasmic space. For the Teneurin ectodomains used in stripe assays, ELISAs and nanobody generation, we use a pHL-Sec vector that contains a TwinStrep tag in the N-terminus (protein sequence: SAWSHPQFEKGGGSGGGSGGSAWSHPQFEK). For cell binding, cell-based aggregation assays and functional assays, full length constructs were used, cloned into the Xho1-Not1 site of the pCAGIG vector (Addgene; Cat#11159)^44^. pCAGIG was modified to express mCherry instead of GFP, for certain experiments, and is then referred to as pCAGIC. Teneurin constructs were cloned to contain an HA tag at the C-terminus (protein sequence: YPYDVPDYA) in the pCAGIG vector, whilst the Latrophilin constructs contained a myc tag at the N-terminus (protein sequence: EQKLISEEDL) in the pCAGIC vector. In addition, the Latrophilin full length constructs contained a PTPmu secretion signal sequenced prior to the myc-tag. We cloned the shRNAs 1 (sequence: GCAGCTCTGGTTGGCATTTAT) and 8 (sequence: GCAGTACATCTTCGAGTTTGA) into the pCAG-miR30 vector (Addgene; Cat#14758). These were designed using the BLOCK-iT^TM^ RNAi Designer tool (https://rnaidesigner.thermofisher.com/rnaiexpress/rnaiDesign.jsp). *amino acid numbering is in A1B1 notation

### Protein expression and purification

For large scale protein expression of the Teneurin ectodomains and biotinylated nanobody, HEK293T cells were grown in supplemented DMEM and 10% FBS in 2125 cm^2^ roller bottles (Grenier Bio-One, Cat#681070). After reduction of FBS to 2%, the cells were transfected with DNA plasmid and polyethylene imine (PEI) (SIGMA-Aldrich; Cat#208727), mixed in a 2-to-1 (PEI to DNA) mg:mg ratio. Cells were left to express the protein for four days and then the media were harvested, clarified by centrifugation and filtered using 0.22 μm sterile filters (Starlab; Cat# S1120-8810). The filtered media was concentrated and buffer-exchanged to diafiltration buffer (1X PBS (Invitrogen; Cat#3002), 20 mM Tris-HCl pH=7.5, 150 mM NaCl). Then this media was passed through a 5 ml HisTrap^TM^ HP column (Cytiva, #Cat17-5248-02) at a flow of 5 ml/min. The column was washed with wash buffer (20 mM Tris-HCl, 300 mM NaCl, 40 mM Imidazole (Sigma-Aldrich, Cat#I3386)) and the protein eluted with 20 mM Tris-HCl, 300 mM NaCl, 500 mM Imidazole. 450 μl of the centre peak fraction was then injected into a pre-equilibrated Superose^TM^ 6 Increase 10/300 GL (Cytiva; Cat#29091598) in 25 mM HEPES-HCl (SIGMA-Aldrich; Cat#7365-45-9)(pH=7.5) and 300 mM NaCl running buffer. Elution fractions were collected, analysed using SDS-PAGE and the peak fractions were either used directly to prepare cryo-EM grids or frozen at –80°C.

For the purification of biotinylated Avi-tagged nanobody, the same protocol was used as before but with some changes. The same DNA:PEI ratio for transfection was used but 90% of the DNA (weight) corresponds to the nanobody-containing vector, and the remaining 10% was BirA-encoding vector. Roller bottles for production of biotinylated protein were supplemented with 0.625 ml of 200 mM biotin after transfection. Purification of the biotinylated nanobody was performed always at 4°C except for size exclusion chromatography. The peak fractions from the HisTrap were injected into a pre-equilibrated Superdex200^TM^ Increase 10/300 GL (Cytiva; Cat# 28990944) in 20 mM Tris-HCl (SIGMA-Aldrich; Cat#7365-45-9)(pH=7.5) and 200 mM NaCl running buffer. Elution fractions were collected, analysed using SDS-PAGE and the peak fractions were either used directly to prepare tetramerised nanobody or frozen at –80°C. For the preparation of the tetramerised NanoTen4, these peak fractions were concentrated and mixed in a 1:8 molar ratio (Streptavidin:Nanobody) with either Streptavidin-Atto647N (Rockland Immunochemicals, Cat#S000-56) or Streptavidin-AlexaFluor^TM^633 (Invitrogen, Cat#S21375), and left at room temperature in the dark for 1 hour. The tetramerised nanobody mixture was then purified using a pre-equilibrated Superdex200^TM^ Increase 10/300 GL (Cytiva; Cat# 28990944) in 20 mM Tris-HCl (SIGMA-Aldrich; Cat#7365-45-9)(pH=7.5) and 200 mM NaCl running buffer. Elution fractions were collected, analysed using SDS-PAGE and the peak fractions were either used directly or frozen at –80°C.

For the N-terminal TwinStrep Teneurin constructs, protein expression was performed in the same way as for their Avi-tagged counterparts. The harvested media was concentrated and buffer-exchanged into diafiltration buffer (1X PBS, 20 mM Tris-HCl pH=8, 150 mM NaCl), passed through a 5 ml StrepTrap^TM^ XT column (Cytiva, #Cat29-4013-17) at a flow of 5 ml/min. The column was washed (50 mM Tris-HCl, 150 mM NaCl, pH=8) and the protein was eluted with elution buffer (50 mM Tris-HCl, 150 mM NaCl, 50 mM biotin). 450 μl of the centre peak fraction was then injected into a pre-equilibrated Superose^TM^ 6 Increase 10/300 GL (Cytiva) in 25 mM HEPES-HCl (pH=7.5) and 300 mM NaCl running buffer. Elution fractions were collected, analysed using SDS-PAGE and the peak fractions were frozen at –80°C.

For the purification of anti-Teneurin4 nanobody used for ELISA analysis and initial characterisation, WK6 bacterial cells (ATCC/Fisher Scientific, Cat# 50-238-2643) were transformed with the desired nanobody construct using heat shock and plated. A single colony was then grown overnight at 37°C in 8 ml of LB medium (Merck, Cat#L3147) with ampicillin. Then 5 ml of the preculture was transferred to 500 ml of complete TB medium (AppliChem, Cat#A0974), incubated 2 hours at 37 °C at 230 rpm until it reached an OD of ∼0.8. The temperature was reduced to 21 °C and incubated for another hour. Protein synthesis was induced with 120 μM (final concentration) of IPTG (Sigma, Cat#I6758-1G) and cells were left to express protein for 16 h at 21 °C and 230 rpm. The cells were pelleted and resuspended with 24 ml ice cold 30 mM Tris pH 7.5, 20% Sucrose, 2 mM EDTA. Cells were incubated on ice for 30 mins and centrifuged for 20 mins at 10000 rpm (fixed angle). The supernatant was saved and the cell pellet resuspended with 25 ml ice cold 30 mM Tris pH 7.5, 5 mM MgSO_4_. Cells were incubated on ice for 20 mins and centrifuged for 20 mins at 10000 rpm. Both supernatants were mixed and the nanobodies purified as explained above for the biotinylated version.

### Cryo-EM grid preparation

Protein samples were used straight after Size Exclusion Chromatography to ensure no protein aggregates deposit onto the grids (A0B1, A0B0, A1B0), or after thawing (A1B1). All samples were cleared by centrifugation prior to grid freezing. For all data presented here, holey carbon Quantifoil® R 1.2/1.3 300 copper mesh grids (Agar Scientific; Cat#AGS143-2) were first plasma cleaned for 2 minutes in a Harrick Plasma Cleaner (Harrick Plasma, USA). 3 μl of protein sample was applied onto the grid and blotted manually with filter paper. The grid was then mounted onto a Vitrobot Mark IV (Thermo Fisher) operating at 4°C (A0B0, A1B0, A0B1) or 21°C (A1B1), 100% humidity. 3 μl of the protein sample was applied again, blotted away with times ranging from 2.5 to 4 seconds before plunge-freezing in liquid ethane.

### Cryo-EM data collection and processing of mTen2 A0B1 dimer

A total of 5,212 movies (Table S1) were recorded using an Talos Arctica cryogenic electron microscope (Thermo Fisher) equipped with a Field Emission Gun (FEG), operating at 200 kV. Micrographs were recorded using a Falcon 4 direct electron detector (Thermo Fisher) at 150,000x magnification (pixel size of 0.94 Å/pix) with an accumulated dose of 40 e-/Å^-2^, and a defocus range of –1.5 to –2.75 μm. All processing was performed in cryoSPARC^101^. Movies were motion corrected using Patch Motion Correction and the CTF estimated using Patch CTF. An initial set of 4,975 particles were picked using Blob Picking, classified in 2D, and the best dimeric classes used as references for Template Picking all micrographs. The resulting 1,037,437 particles were extracted with a box size of 658 Å^2^ and binned by 2. Junk particles were removed through successive rounds of 2D classification. Two ab initio models were then generated with 221,220 “good”, unbinned particles, and further 3D classified using the Heterogeneous Refinement job. The best 3D class containing 116,348 particles was further refined using Non-Uniform Refinement^102^, while optimising per-particle defocus and per-group CTF parameters and with C2 symmetry applied. This yielded a final map of 3.48 Å (FSC = 0.143, as reported by cryoSPARC). Details can be found in Extended Data Fig. 1A.

### CryoEM data collection and processing of mTen2 A1B1, A1B0 and A0B0 dimers

Datasets for mTen2 A1B1, A1B0 and A0B0 dimers were all collected on a Titan Krios electron microscope (Thermo Fischer) using a nominal magnification of 105,000x (0.83Å/pix) and operating at 300 kV. Movies were recorded using a K3 Summit direct electron detector with a Bioquantum energy filter (Gatan) (20 eV slit), an accumulated dose ranging between 41.5 and 42.3 e-/Å^-2^, and a defocus range of –1 to –2 μm. A total of 8,975, 5,136 and 6,108 movies were recorded for mTen2 A1B1, A1B0 and A0B0 respectively (Table S1). For all three datasets, processing was carried out in cryoSPARC ^101^. All movies were first motion corrected using Patch Motion Correction followed by CTF estimation with Patch CTF. 2D references generated either from previous pilot datasets or a subset of Blob Picked particles were used for Template Picking. 6,799,146, 3,672,845 and 4,649,892 particles were initially extracted for A1B1, A1B0 and A0B0 datasets respectively, all with a box size of 301 Å^2^, and junk particles discarded through multiple rounds of 2D classification. Three ab initio models were generated for each dataset with “cleaned” particles, which were further 3D classified through one or two rounds of Heterogeneous Refinement. The best 3D class contained 128,786, 167,798 and 260,530 particles for A1B1, A1B0 and A0B0 respectively. Those were further refined using Non-Uniform Refinement ^68^, while optimising per-particle defocus and per-group CTF parameters as well as with C2 symmetry applied, yielding maps of 2.54 Å, 2.55 Å and 2.54 Å respectively. For A1B1 and A0B0 datasets, particles contributing to each refined map were further sorted via 3D Classification into 10 classes. In each case, the class that had both NHL domains resolved was then refined using Non-Uniform Refinement ^102^, resulting in a final resolution of 2.80 Å and 2.82 Å for A1B1 and A0B0 maps respectively (FSC = 0.143, as reported by cryoSPARC). Details can be found in Extended Data Fig. 1A.

### Molecular dynamics simulations and hydrogen bond identification

To perform an all-atom molecular simulation of Teneurin, we followed the approach outlined by Lemkul^103^ that we used earlier to study the GPC3-Unc5 receptor complex^42,103^. Briefly, we built a simulation model of the Teneurin homodimer based on the structure obtained by cryo-electron microscopy (Ten2-Ten2 A1B1 dimer). We removed all glycosylations from the structure and kept the disulfide bridges. We then used the Amber ff14SB force field to model this structure. Because the Amber ff14SB force field lacked an improper dihedral interaction for the HisD protonation state (6 His residues out of a total of 82), we manually set the protonation state of all His residues to HisE. We embedded the protein in a cubic box with edges at least 2.0 nm away from the protein. We solvated the protein in TIP3P water, neutralized the system and added 150 mM NaCl. The resulting system had a box size of (22.4 nm)^3^ and contained 1098381 atoms. We have used GROMACS 2023.2^104^ to perform the simulations. During the entire simulations we imposed position restraints on all backbone atoms (C, C_α_ and N), but not on the side chains, to preserve the overall protein structure. We minimized the system using the steepest descent method, then carried out a 100 ps NVT equilibration followed by a 100 ps NPT equilibration and a 1 µs production run. In all these simulations we used a Éme step of 2 fs, we set the temperature at 310 K using the V-rescale thermostat^105^ with a Éme constant ι_T_ of 0.1 ps and we set the pressure at 1 bar using the Parrinello-Rahman barostat^106^ with a time constant of 2.0 ps and an isothermal compressibility of 4.5 × 10^-5^ bar^-1^. We employed the MDAnalysis toolkit^107,108^ to analyse MD simulation trajectories. For contact analysis, we have used the Distance_array function between the centres of mass of residues with a cutoff of 8 Å. To identify hydrogen bonds between the Teneurin monomers, the HydrogenBondAnalysis tool^109^ provided in using a donor-acceptor distance cut-off of 3.0 Å and a cut-off angle of 150°. We used the aforementioned protocol but without constraints on the backbone for 1 μs.

### Stipe assays and immunostaining

We prepared the stripe assays as previously described^3^. 50 μg/ml of Fc recombinant protein (Jackson Immunoresearch; Cat#009-000-008; AB_2337046), Ten4 WT, Ten4 nT or Ten4 nL, were mixed with AlexaFluor^TM^ 647-conjugated (Thermo Fischer Scientific (Cat#A-11014; AB_1500628) anti-hFc antibody (Thermo Fisher Scientific; Cat#62-8400; AB_2337530) in PBS (Life Technologies; Cat#10010023). Proteins were injected into matrices (90 μm width) (17546017) and placed on 60 mm dishes, resulting in red fluorescent stripes. After 30 min incubation at 37°C, dishes were washed with PBS and matrices removed. Dishes were coated with 50 μg/ml Fc or Ten4WT/nT/nL protein mixed with 150 μg/ml anti-hFc (Jackson ImmunoResearch, Cat#62-8400) for 30 min at 37°C and washed with PBS. Stripes were further coated with 20 μg/ml Laminin in PBS overnight and washed with PBS next morning. Cortical neurons (E14.5) were cultured on the stripes in Neurobasal medium supplemented with B27. After 24 hours neurons were fixed with 4% PFA in PBS for 10 min at room temperature (RT) and stained as explained below. The numbers of ISH-positive pixels on red or black stripes were quantified with ImageJ (version 2.9.0/1.53t)^110^ after sorting on low/mid/high Ten4 expressing cells using Cellprofiler ^111^. For both software programmes, we used custom-made automated macros, which are available upon request.

### RNA In situ hybridization (ISH) in cultured neurons

Cultured neurons on stripes were fixed in 4% PFA for 10 min and dehydrated with increasing ethanol concentrations solutions (50, 70, 100%) in PBS for 5min at room temperature.

Stripes were kept in 100% ethanol at –20°C until further use. Stripes were pre-treated using the RNAscope Universal Pretreatment Kit (Advanced Cell Diagnostics; Cat#322380). RNA In Situ Hybridizations (ISH) were performed using the RNAscope Fluorescent Multiplex Reagent Kit (Advanced Cell Diagnostics; Cat#323100) according to manufacturer’s instructions. The target gene (Ten4) was detected with the probe: Mm-Tenm4-C1 (RNAscope; Cat#555491). Following ISH, nuclei were counterstained with DAPI before mounting. Images were acquired using a Zeiss LSM880 confocal laser scanning microscope or a THUNDER imager (Leica) using a 10x objective and processed with ImageJ software.

### In utero electroporation

In utero electroporation was performed at E13.5 with anesthetized C57BL/6 mice as previously described^3^. DNA plasmids were used at 2 µg/µl and mixed with 1% fast green (Sigma-Aldrich, final concentration 0.2%). Plasmids were injected into the ventricle with a pump-controlled micropipette. After injection, six 50 ms electric pulses were generated with electrodes confronting the uterus above the ventricle. The abdominal wall and skin were sewed, and the mice were kept until E16.5 embryonic stage. To knockdown Ten4 by shRNA in vivo, we used two shRNAs embedded in the in the pCAG-miR30 vector, with the following sequence: shRNA#1 (sequence: GCAGCTCTGGTTGGCATTTAT) and #8 (sequence: GCAGTACATCTTCGAGTTTGA). This shRNA was validated in HEK293T cells, by co-transfection with Ten4 followed by Western Blotting (see below). For the CRISPR-Cas9 editing of the endogenous Ten4 locus, we used the Alt-R HDR Design tool on the IDT website.

Oligonucleotides and Cas9 were ordered from Integrated DNA technologies (IDT) using the Alt-R design tools and reagents. This includes the transactivating CRISPR RNA (IDT #:238355667), scrambled negative control crRNA (IDT #238355668 and #238355669), and the targeted CRISPR RNA (crRNA) sequences. Ten4 nT mutant was generated by targeting Exon30 (position: 96892998) with the following sequence: ACCTGACCGGCGTGAACGTGACA. For Ten4 nL, we targeted Exon29 (position:96888889) with the following sequence: GTGTTCGGCAGAGACCTGAGA, and Exon 30 (position: 96892922) with: GACGCCCACAGGAAGTTCACCCTGAGGATCCTGTAC. We followed the IDT protocol, where tracrRNA (IDT #238355667) and crRNA (400 µM stock) are hybridized in a 1:1 molar solution at 95°C for 5min and held at room temperature for 10min, resulting in the final guide RNA (gRNA). The final solution was composed of 1.5 µl of gRNA (200 µM Stock), 2ul of HDR template (200uM), 1 µl CAS9 (IDT #1081058), 1µl pCAGIG (0.6ug/l final concentration), 1 µl Fast Green (final 0.2%) to a final volume of 10 µl. This solution was incubated at 37°C for 10 min before surgery.

### Mass spectrometry of dissected cortical layers

Fresh E15.5 mouse brains (N=12 brains) were divided into 4 replicate groups and each replicate was then manually dissected in cold PBS under and Olympus SZX10 stereomicroscope to isolate the different cortical layers: VZ/SVZ, IZ and CP. Each sample for each replicate group was homogenized separately for 1 min at 4°C with an electric homogenizer in 50 μl of the following lysis buffer: 50 mM Tris-HCL (pH 7.4), 150 mM NaCl, 2 mM EDTA, 1% Triton X-100 and protease inhibitors (Roche, Cat#04693116001). Samples were incubated on ice for 20 min and centrifuged for 10 min at 3000 rpm. The supernatant was collected, and then samples were processed for mass spectrometry and analysed (MaxQuant run, Proteomic facility, Max Planck Institute of Biological Intelligence, Martinsried, Germany). All reported readings for protein readings were normalized against PGK (house-keeping gene).

### Shadow imaging

Three electroporated brains, coming from different animals, per condition were dissected and fixed in 4% PFA overnight, then washed and stored in PBS. We processed the brains with a vibratome to produce 100 µm sections. These brain slices were incubated at 4°C for 2.5 h and then at R/T for 1.5 h in PBS with 5 µg/mL Wheat Germ Agglutinin, CF®594 Conjugate (Biotium, Cat#29064-1). Then, they were washed three times with PBS for 30 minutes. Finally, they were mounted in Fluoromount-G Mounting Medium (Southernbiotech, Cat#0100-01) on a glass side under a cover slip. For the cortical plate, a standard research confocal microscope was used (Zeiss LSM880) with a 63X oil immersion objective (NA 1.4; Plan-Apochromat 63X/1.4 Oil DIC M27). Image acquisition was controlled by the microscope’s commercial software (ZEN 2.3 sp1). Fluorophores were selectively excited by different lasers at wavelengths of 488 nm for GFP (mutant neurons), 561 nm for extracellular label (shadow imaging), and 633 nm for BLBP immunofluorescence. Image stacks were acquired with a x-y pixel size of 220 nm, a z-step of 1.5 μm and pixel dwell time of 2.05 μs. The imaging of the intermediate zone was done using another but similar confocal microscope (Leica DMI6000 TCS SP8 X) also equipped with a 63X oil immersion objective of the same numerical aperture (NA 1.4; HC PL APO CS2 63x/1.40 OIL). Image acquisition was controlled by the microscope’s commercial software (LAS X). Fluorophores were selectively excited by different lasers at wavelengths of 488 nm for GFP (mutant neurons), 594 nm for extracellular label (shadow imaging), and 647 nm for BLBP immunofluorescence. Image stacks were acquired with a x-y pixel size of 361 nm, a z-step of 1.5 μm and pixel dwell time of 3.16 μs.

Areas that contained GFP-positive and GFP-negative cells alongside BLBP signals were selected for analysis from three separate slices coming from different animals as independent replicates. To obtain a measure of cell size, we outlined the somata of GFP-positive (mutant) and GFP-negative (control) neurons and expressed the enclosed area in square micrometres (mm^2^). To assess the association between neurons (mutant and wildtype control) and RGC fibres labelled by immunofluorescence (antibody directed against BLBP), the images were thresholded using the threshold function in ImageJ. The threshold was consistently set to 1%, so only the most intense 1% of pixels were considered as part of our object of interest. These resulting areas served as a local measure of the RGC fibres.

Using ImageJ’s automatic ROI enlargement function, we expanded the outlined somata of the cells by 2 μm. Then, we measured the overlap between the RG fibres and the expanded somata using the ImageJ ROI Manager. To separate the neurons depending on low, mid, and high levels of RGC fibre attachment, we used the following intervals: 0-3.5 μm^2^ for low, 3.5-6.5 μm^2^ for mid, and 6.5-10 μm^2^ for high.

### Cell-based Binding Assay

HEK293T cells grown on coverslips were transfected using HA-tagged constructs in pCAGIG vector with 3 μg of DNA and 9 μl of PEI. Eighteen hours after transfections, cells were incubated with buffer (HBSS (Life Technologies; Cat#14170088) with 1% BSA and 10 mM HEPES (pH 7.5)) for 30 minutes on ice, and then with buffer containing 0.5 μg purified His-tagged protein per coverslip that was previously pre-clustered (30 mins at room temperature) with anti-His (mouse; Thermo Fisher Scientific; Cat#372900; RRID: AB_2533309) in a 1:2 (protein:antibody) ratio (mass:mass) for 60 minutes on ice. Cells were then washed with PBS and fixed with 4% PFA for 20 minutes, and then washed using PBS supplemented with 50 mM ammonium chloride (SIGMA-Aldrich; Cat#326372-100G). Cells were then incubated in the dark with anti-mouse-Cy3 (Invitrogen; Cat#A10521; AB_10373848) in a 1:7.5 (protein:antibody) ratio (mass:mass) in buffer for 60 minutes on ice. The cells were washed with PBS, stained with DAPI (0.1 μg/ml), washed with PBS and mQ water, and then mounted using Immu-Mount (Fischer Scientific; Cat#10622689) into SUPERFROST microscope slides (Fischer Scientific; Cat#12372098). For the *in vitro* validation of NanoTen4, the same protocol was used but NanoTen4 was added to the cells for 120 minutes on ice and there was no secondary antibody step. Imaging for data analysis was done with a Nikon ECLIPSE TE2000-U inverted fluorescence microscope or a THUNDER Imager (Leica). Analysis was performed in ImageJ (version 1.54f)^110^ using the co-localisation macro used previously^33^. The co-localisation area of red pixels (soluble bound protein) with green pixels (transfected cells) was normalised against the area of the transfected cells and converted into a percentage. For the statistical analysis we used a one-way ANOVA test, with a Tukey’s post-hoc test in Graphpad Prism (version 10 for MacOS, GraphPad Software, San Diego, California USA, www.graphpad.com). Significance was determined when p < 0.05. Pictures showed in Figures were imaged using a THUNDER Imager (Leica).

## shRNA *in vitro* validation

HEK293T were cultured in 6-well plates on coverslips. 24 hours after culturing, the cells were co-transfected with shRNA (both #1 and 8 constructs) and full length Ten4 A1B1 in a 3:1 μg ratio with Fugene (Promega; Cat#E2693) (1:3 (v/m) ratio with total DNA). Cells were harvested 72 hours after transfection, sonicated (2 pulses of 5 and 7 seconds respectively on ice). Samples were loaded into a 3-8% Tris-Acetate gels (Thermo Fischer Scientific; Cat#EA03785BOX), run for 15 mins at 150 V and then for 40 mins at 180 V. The gels were transferred in the NuPAGE^®^ XCellII^TM^ Blot transfer module for 90 minutes at 30 V onto a nitrocellulose membrane (Cytiva, Cat#10600013), sandwiched with two blotting papers and five sponges, submerged in 1X NuPAGE^®^ Transfer Buffer (Invitrogen^TM^, Cat#NP00061) supplemented with 20% ethanol. To confirm successful transfer, the HiMark^TM^ Pre-stained Protein ladder (Invitrogen, Cat#LC5699) was used. After transferring, the membrane was blocked for 30 minutes using 3% BSA (w/v) (Invitrogen^TM^, Cat#A7906) in PBST (1X PBS with 0.1.% Tween20) and washed with PBST only for 30 mins. Then the membranes were incubated in the primary antibody (mouse anti-HA (Sigma-Aldrich, Cat#H3663) prepared in PBST with 3% BSA in a 1:1000 dilution) solution for 1 hour. After incubation, the membrane was washed for 30 minutes with PBST and incubated for 1 hour with the secondary antibody solution (anti-Mouse IgG HRP (Sigma-Aldrich, Cat#A0168) diluted 1:10000 in PBST with 3% BSA). Then the membrane was washed with PBST for 45 minutes and developed with the ECL^TM^ Western blotting detection reagents (Cytiva, Cat#RPN2106) and visualised using Amersham^TM^ Hyperfilm^TM^ ECL^TM^ films (Cytiva, Cat#28-9069-35). The Western Blots were quantified using ImageJ^110^ using the intensity of the construct band normalised by subtracting against the intensity of the non-transfected band for control purposes. For the statistical analysis we used a one-way ANOVA test, with a Tukey’s post-hoc test in GraphPad Prism (version 10 for MacOS, GraphPad Software, San Diego, California USA, https://www.graphpad.com/). Significance was determined when p < 0.05.

### Cell-cell aggregation assay

K562 suspension cells were cultured in RPMI-1640 media (no phenol red) (Invitrogen; Cat#11835030) supplemented with 10% FBS and 5% L-Glutamine. The cells were harvested by a 3 min spin at 200g, washed with PBS, spined again and resuspended in R buffer (Neon Transfection System 100 μL Kit; Thermo Fischer Scientific; Cat#MPK10025). Cells at a concentration of 2×10^7^ cells/ml were transfected with control pCAGIG/pCAGIC plasmids, or those coding for mTen2,4 or mADGRL1-3 constructs using the Neon transfection system for electroporation (Settings: 1450V, 3 pulses, 10 ms). Eighteen hours after transfection, cells were harvested and used at a concentration of either 2 x 10^5^ cells/ml or 4 x 10^5^ cells/ml in aggregation media (Neurobasal-A media without phenol red (Thermo Fischer Scientific; Cat#12349015) supplemented with 2 mM L-glutamine (Life Technologies; Cat#25030-024), 10% FBS, 4% B-27 and 20 mM HEPES) in a 24-well plate. Cells were then left to aggregate at 37°C, 5% CO_2_ and 250 rpm for 90 minutes. After the incubation, the cells were imaged directly in the 24-well plate after a slight shake using a Nikon ECLIPSE TE2000-U inverted fluorescence microscope. The total area of cells and the total area of the aggregates for each picture were calculated using the Analyze particle tool in ImageJ ^110^. The threshold used to distinguish cells and aggregates was determined at 1284 μm^2^ (>3/4 cells). For the statistical analysis we used a one-way ANOVA test, with a Tukey’s post-hoc test in GraphPad Prism (version 10 for MacOS, GraphPad Software, San Diego, California USA, www.graphpad.com). Significance was determined when p < 0.05. Pictures showed in Figure 2 and Extended Data Fig. 2 were imaged using a THUNDER Imager (Leica).

### Cell surface expression in K562 cells

For surface staining, K562 cells were harvested eighteen hours after electroporation (performed as explained above) and cooled to 4°C for 30 mins. Cells were spined at 200g for 3 mins (4°C) and the media aspirated. This process of spinning and aspirating was performed in between all following steps. All the steps in this protocol are performed on non-permeabilised cells, so that the antibody only detects protein that has been trafficked to the cell surface where the tag is exposed to the antibodies. We also perform all steps at 4°C to abolish endocytosis. The cells were then incubated on ice with blocking buffer: HBSS with 1% BSA and 10 mM HEPES (pH 7.5) for 30 minutes on a shaker. For Ten2,4 expressing cells, cells were incubated for 1 hour on ice on a shaker. We used anti-HA (mouse; SIGMA-Aldrich; Cat#H3663; RRID: AB_262051) antibody that was pre-clustered with secondary anti-Mouse antibody-Cy3 (Invitrogen; Cat#A10521; AB_10373848) for 40 minutes (ratio 1:7.5 for primary:secondary antibody) at room temperature in the dark. Cells were washed with PBS, fixed with 4% PFA for 20 minutes, washed with PBS with ammonium chloride and stained with DAPI (0.1 μg/ml) for five minutes on ice and shaking. Cells were then resuspended on 20 μl

Immu-Mount and put on the centre of the slide. Finally, a rectangular coverslip was put on top of the resuspended cells. The cells were imaged on a using a Nikon ECLIPSE TE2000-U inverted fluorescence microscope (20X magnification). Colocalization for the surface quantification was performed using ImageJ ^110^ with the colocalisation tool and a macro that can be made available upon request. This colocalisation analysis compares the pixels in both channels (green (transfected cells) and red (anti-HA/Cy3 clustered antibodies)) and determines which pixels have a signal in both channels. A ratio of 10% between red and green channels was used. As the cells are non-permeabilised, the signal coming from the red channel (Cy3) can only come from the HA-tagged proteins on the cell surface, if present. The area of the pixels that are colocalised between both channels is the normalised by the total transfected cell area. For the statistical analysis we used a one-way ANOVA test, with a Tukey’s post-hoc test in GraphPad Prism (version 10 for MacOS, GraphPad Software, San Diego, California USA, www.graphpad.com). Significance was determined when p < 0.05.

### Cell surface expression in HEK293T cells

HEK293T cells grown on coverslips were transfected (24 hours after seeding, at a confluence of 80%) using full length Teneurin constructs in pCAGIG vector with 3 μg of DNA and 9 μl of PEI. All the steps in this protocol are performed on non-permeabilised cells, so that the antibody only detects protein that has been trafficked to the cell surface where the tag is exposed to the antibodies. We also perform all steps at 4°C to abolish endocytosis. Eighteen hours post-transfection, cells were incubated with buffer (HBSS with 1% BSA and 10 mM HEPES (pH=7.5)) for 30 minutes on ice and then with buffer containing 1 μg of mouse anti-HA antibody for 1 hour. Cells were then washed with PBS and fixed with 4% PFA (Sigma-Aldrich, Cat#158127-100G) for 20 minutes and washed again with PBS supplemented with 50 mM ammonium chloride. Cells were then incubated in the dark with anti-Mouse Cy3 (Abcam, Cat#ab97035; RRID: AB_10680176) in a 1:7.5 primary:secondary ratio in buffer for 1 hour. The cells were then washed with PBS, stained with DAPI (0.1 μg/ml) and mounted using Immu-Mount (Thermo Fisher Scientific, Cat10622689). Imaging was performed using a Nikon ECLIPSE TE2000-U inverted fluorescence microscope. Quantification was performed using ImageJ ^110^ using the with the colocalisation tool and a macro that can be made available upon request. This colocalisation analysis compares the pixels in both channels (green (transfected cells) and red (anti-His/Cy3 clustered antibodies)) and determines which pixels have a signal in both channels. A ratio of 10% between red and green channels was used. As the cells are non-permeabilised, the signal coming from the red channel (Cy3) can only come from the soluble His-tagged proteins on the cell surface, if present. The co-localisation area of green pixels (transfected cells) with red pixels (cell-bound protein) was normalised against the area of transfected cells and converted into a percentage. For the statistical analysis a one-way ANOVA test was used, with a Tukey’s post-hoc test in GraphPad Prism (version 10 for MacOS, GraphPad Software, San Diego, California USA, https://www.graphpad.com/). Significance was determined when p < 0.05.

### Surface biotinylation of surface expressed proteins

HEK293T cells were grown on T75 flasks until 90% confluent. Then cells were transfected using 780μl of optiMEM, 70.2 μl of PEI, and 23.4 μg of DNA (empty pCAGIG, Ten4 WT, Ten4 nT or Ten4 nL). To assess protein expression on K562, cells were electroporated as explained above (see cell-cell aggregation assay). Both cell lines were incubated overnight (∼16 hours) at 37°C. To perform the surface biotinylation and extraction of surface-bound proteins, we used the Pierce™ Cell Surface Biotinylation and Isolation Kit (Cat#A44390) and followed manufacturer’s guidelines. For HEK293T cells, we removed the media and washed cells with 10 mL of ambient PBS per T75. We dissolved the contents of one vial of Sulfo-NHS-SS-Biotin in 24 mL of ambient PBS. We added 5 mL of the biotin solution to one T75 and incubated the T75s for 10 minutes at room temperature. After the incubation, we removed labelling solution and washed cells twice with 10 mL ice-cold TBS. removing the TBS after each wash. We scrapped the cells into 10 mL ice-cold TBS and put them on a 50 mL conical tube on ice. We rinsed the scraped T75s with 5 mL each of TBS and added the rinse volume to transferred cells. We then centrifuged cells at 500 × *g* for 3 minutes at 4°C and discarded the supernatant. For K562 cells, we grew the cells in suspension to ∼4 × 10^5^ cells/ml having a total of 15 ml.

We harvested and centrifuged cells at 300 × *g* for 3 minutes and discarded the media. We washed the cells with 10 ml of PBS, centrifuged them at 300 × *g* for 3 minutes and discarded the wash. We added 5 ml of the biotin solution (prepared as explained above) to the tube and resuspended the cells. These were incubated for 10 minutes at room temperature. We then centrifuged the cells at 300 × *g* for 3 minutes and removed label solution. Cells were washed with 10 ml ice-cold TBS two times with a centrifugation step in between at 300 × *g* for 3 minutes discarding the supernatant. For both cell lines, we added 50 μl of 10X cOmplete^™^, Mini, EDTA-free Protease Inhibitor Cocktail (Sigma Aldrich, Cat# 11836170001) to 500 μl of Lysis Buffer and added 250 μl to the cell pellets. We resuspended the cells 20 times and transferred the cells to Eppendorf tubes. These were vortexed for 5 seconds before and after an incubation on ice for 30 mins. The cell lysate was centrifuged at 15,000 × *g* for 5 minutes at 4°C. The supernatant was incubated with 125 μl of NeutrAvidin^TM^ Agarose slurry for 30 minutes at RT on columns. After incubation, the column was centrifuged for 1 minute at 1,000 × *g*. The columns were then washed 4 times with the wash buffer. We mixed 225 μL Elution Buffer and 25 μL DTT stock solution and added 100 μL of the buffer to the resin. This reaction was incubated for 30 minutes at room temperature.

The supernatant was then collected after centrifuging for 2 minutes at 1,000 × *g*. The eluate was then analysed using SDS-PAGE and Western Blot as explained above (see shRNA validation *in vitro*). We used mouse anti-HA (Sigma-Aldrich, Cat#H3663) and anti-mouse HRP (Sigma-Aldrich, Cat#A0168) to detect Ten4 (WT/nT/nL) protein levels. As a loading control, we used rabbit anti-Transferrin Receptor (Abcam, Cat#ab214039) and anti-rabbit HRP (Abcam, Cat#ab6721) to detect the levels of extracted Transferrin Receptor.

### Liquid chromatography-mass spectrometry (LC-MS) and mass spectrometry analysis

For sample preparation, 17.5 µg protein present in 20 mM Tris, 300 mM NaCl buffer, were treated with 500 U of PNGaseF (NEB P0705S) for 1 h at 37 °C. Subsequently, the samples were denatured in 4 M Urea/0.1 M ammonium bicarbonate buffer pH 8.0, followed by 30 min incubation with 10 mM TCEP at room temperature and 30 min incubation with 50 mM 2-chloroacetamide at room temperature in the dark. Samples were diluted 4 times in 0.1 M ammonium bicarbonate buffer and digested with protease chymotrypsin (1 µg enzyme: 40 µg protein), 10 mM calcium chloride, overnight at room temperature. Digestion was stopped with the addition of formic acid (5 %). Digested peptides were centrifuged for 30 min at 12,700 rpm at 4 °C. The supernatant was loaded onto C18 stage tips, pre-activated with 100 % acetonitrile and 0.1 % formic acid. Centrifugation was carried out at 4000 rpm at room temperature. Next digested peptides were loaded onto the C18 column. Subsequently, peptides were washed with 0.1 % formic acid and eluted in 50 % acetonitrile / 0.1 % formic acid. Eluted peptides were dried in a speed-vac before resuspension into 5% acetonitrile / 5 % formic acid for LC-MS/MS analysis. Peptides were separated by nano liquid chromatography (Thermo Scientific Easy-nLC 1000) coupled in line a Q Exactive mass spectrometer equipped with an Easy-Spray source (Thermo Fisher Scientific). Peptides were trapped onto a C18 PepMac100 precolumn (300µm i.d.x5mm, 100Å, Thermo Fisher Scientific) using Solvent A (0.1% Formic acid, HPLC grade water). The peptides were further separated onto an Easy-Spray RSLC C18 column (75um i.d., 50cm length, Thermo Fischer Scientific) using a 30 minutes linear gradient (15% to 38% solvent B (0.1% formic acid in acetonitrile)) at a flow rate 200nl/min. The raw data were acquired on the mass spectrometer in a data-dependent acquisition mode (DDA). Full-scan MS spectra were acquired in the Orbitrap (Scan range 350-1500m/z, resolution 70,000; AGC target, 3e6, maximum injection time, 50ms). The 10 most intense peaks were selected for higher-energy collision dissociation (HCD) fragmentation at 30% of normalized collision energy. HCD spectra were acquired in the Orbitrap at resolution 17,500, AGC target 5e4, maximum injection time 120ms with fixed mass at 180m/z. Charge exclusion was selected for unassigned and 1+ ions. The dynamic exclusion was set to 5s. For data processing, tandem mass (MS/MS) spectra were searched using PEAKS X software version 10.0 as follows against a protein sequence database containing 20,606 protein entries, including 20,323 *Homo Sapiens* proteins (Uniprot release from 2024-02-16), 283 common contaminants and mouse Teneurin 2 and 4 WT and variants sequences. During database searching for full chymotryptic peptides, cysteines (C) were considered to be fully carbamidomethylated (+57,02, statically added), methionine (M) to be fully oxidised (+15,99, dynamically added), all N-terminal residues to be acetylated (+42,01, dynamically added) and asparagine to be converted into aspartic acid (−0.98, dynamically added). Three missed cleavages were permitted. Peptide mass tolerance was set at 20 ppm on the precursor and 0.6 Da on the fragment ions. Data was filtered at a False Discovery Rate (FDR) below 1% at the peptide level. The mass spectrometry proteomics data have been deposited to the ProteomeXchange Consortium via the PRIDE partner repository with the dataset identifier PXD058769.

### Enzyme-linked Immunosorbent Assays (ELISA)

NuncTM Maxisorp^TM^ 96-well plates (Thermo Scientific, Cat#735-0083) were coated with TwinStrep Teneurin2-4 or Latrophilin3 at a concentration of either 25 nM or 15 nM and left overnight at 4°C. The plates were washed three times with PBS supplemented with Tween20 (PBST) and dried. All the wells were blocked with BlockerTM Casein in PBS (ThermoScientific, Cat#37528) for 30 minutes at room temperature. After blocking, plates were washed with PBST three times and 50 μl of the anti-Ten4 nanobodies were added at a concentration of 500 nM diluted in Casein. We used an anti-GPC3 nanobody (Nano^glue^) ^42^ as a negative control. Blank wells were left in Casein in PBS. Plates were incubated for 60 minutes at room temperature and then washed 3 times with PBST. 50 μl of the primary antibody solution was then added to the wells and left for 1 hour at room temperature. For the nanobody-containing wells, mouse anti-His antibody (Thermo Fisher Scientific; Cat#372900; RRID: AB_2533309) at a dilution of 1:1000 was used, but to create a standard curve, we used a titration of mouse anti-Strep antibody (IBA, Cat#2-1507-001) starting from 5 μg/ml was added to wells that were coated with Ten4 but were not added nanobody. The plates were all washed with PBST and all wells were added 50 μl of a solution containing anti-mouse IgG alkaline phosphatase antibody (Sigma-Aldrich, Cat#A1418-.25ML, 1:10000 in casein) and incubated for 30 mins at room temperature in the dark. Plates were washed with PBST and PBS and revealed with 100 μl of a buffer containing 1x diethanolamine buffer (Fischer, Cat#34064) and 20 mg of 4-nitrophenylphosphate tablet (Sigma, Cat#N2765-50TAB) diluted to a concentration of 1mg/ml. Plates were then read every one minute noting the 405 nm absorbance until one sample reached an OD of 1.3. ELISA readings were standardised by calculating the linear fit for the Streptavidin standards and applying the linear equation to the nanobody readings to calculate the normalised response. We reported the results as arbitrary units (AU). For the Kd calculation, a titration of anti-Ten4 nanobody was prepared starting from 10000 nM and readings were produced following the same protocol. After standardisation of the readings, the Kd and fit were calculated using an equation described before ^112^.

### Stimulated Emission Depletion (STED) microscopy

A shRNA control (miR30 control GFP+) slice was immunostained following a similar protocol described above. The brain slice (thickness 75 μm) was incubated for 2 hours at room temperature in PBS with 0.5% Triton X-100 (Sigma-Aldrich, Cat#X100-100ml) and then rinsed with PBS. The slice was incubated overnight at 4°C in NanoTen4 and anti-BLBP antibody (Millipore-Merck, Cat#Abn14), both diluted 1:50 in PBS with 0.3% Triton X-100, 0.2% BSA, 0.2% Gly, 0.2% Lys and 5% donkey serum. The slice was washed three times with PBS and further incubated three hours at 4° with a dilution of 1:250 of anti-rabbit AlexaFluor 594 secondary antibody (Fisher Scientific, Cat#A21207). The slice was washed in PBS for 30 minutes and mounted using Mowiol (pH 8.5, no propyl gallate) and a #1.5H high precision coverslip (Electron Microscopy Sciences, Cat#71861-054-C).

For super-resolution STED imaging, the sample was acquired with a STELLARIS 5 TauSTED Xtend confocal microscope system (Leica Microsystems) using a HC PL APO 100x/1.40 OIL STED WHITE oil objective. NanoTen4 and anti-BLBP were imaged in 2D TauSTED mode and GFP signal in conventional confocal mode. Z-stacks of ∼10-15 μm hight were acquired with 500 nm z-steps and a pixel size set to 21-23 nm. A white light laser set to 100% power was used for excitation and a 775 nm solid state laser was used for depletion of both STED channels. Channel shifts between STED and confocal channels were determined from acquisitions of 100 nm TetraSpek beads (ThermoFisher) and corrected using ImageJ ^110^ (12 pixels to the left and 8 pixels up).

The STED images were analysed using two custom macros on ImageJ ^110^ (available upon request). ROIs were first generated to select the RGC fibres using the BLBP STED images. Then an area around the fibre at a maximum distance of 1.5 μm from the centre of the fibre was selected in an unbiased manner, independent of the NanoTen4 staining. NanoTen4 STED images were first denoised by applying a median (0) filter and intensity-normalised before converting to a binary image applying an arbitrary threshold of 1000. Next, the area of NanoTen4-positive pixels within the ROIs generated in the previous step for all stacks and images was determined and the enrichment of NanoTen4 for each individual RGC fibre calculated relative to the surrounding area for each image. Averaged values for each image in the stacks (individual image) were reported, and the ratios between the enrichment of RGC fibre and the surrounding area were determined. For the classification of stacks into high/mid/low NanoTen4 stained areas, we calculated the thresholds for each level by using descriptive statistics and using the values for the 25 and 75 percentiles for the enrichment of NanoTen4 on RGC fibres on the cortical plate as thresholds.

### Basescope^TM^ and sequencing validation of CRISPR-introduced nT and nL mutations

BaseScope^TM^ assays were performed following guidelines (BaseScope™ Reagent Kit-v2 RED) provided by the supplier (Advanced Cell Diagnostics, Newark, CA). In brief, electroporated CRISPR brains were fixed in 4% PFA overnight and cut using a cryostat to a 10 mm thickness. 10 mm Cryo-sections were pretreated with hydrogen peroxide, target retrieval and protease III. Then BaseScope^TM^ probes (the three probes were all purchased from Advanced Cell Diagnostics and sequences are available in the Key Resources table) were applied for 2 hours at 40°C in a HybEZ oven follow by the amplificator reagents following the protocol indications. Finally, slides were incubated with Fast Red for 10 min at room temperature in the dark. Then, nuclei were counterstained with DAPI, and sections were mounted with Mowiol. Images were acquired using Thunder microscope (Leica) with a 20x objective and using 2 cameras, the colour for Basescope detection and the fluorescent camera for GFP/DAPI detection. Images were processed with ImageJ ^110^ and Cell profiler software ^111^.

For sequencing, electroporated CRISPR brains were dissected in cold HBSS using a Leica MZ10 Stereomicroscope equipped with a fluorescent lamp. The electroporated region from each brain was collected into an Eppendorf tube, centrifuged, and the tissue was stored at –80 °C until further processing. DNA was extracted by boiling the samples in 50 µl of 50 mM NaOH at 95 °C for 5 minutes, followed by neutralization with 5 µl of 1.5 M Tris-HCl (pH 8.8). PCR was used to amplify the targeted region for the Ten4 TT mutation (Exon 30, position 96892998) using the following primers: Forward: GATCTACGATGACCATCGCA and Reverse: TGCAAAGATCCGGGATGTG. The PCR reaction was performed using Invitrogen™ Platinum™ II Hot-Start PCR Master Mix (2X), and the amplicons (220 bp for nT and 157 bp for nL) were purified using the QIAquick PCR Purification Kit. The quality assessment, sequencing, and raw data control for each sample were performed using the INVIEW CRISPR Check (150–270 bp) service from Eurofins (https://eurofinsgenomics.eu/en/next-generation-sequencing/ngs-built-for-you/inview-crispr-check/). The average quality score ranged from 88.59% to 97.20%, and the total number of read pairs was approximately 5 million per sample. We used R, along with the ShortRead and ggplot2 libraries, to identify the correctly CRISPR-edited transcripts: ACCTGACCGGCGTGAACGTGACA for the nT mutation and GTGACAGTGTTCGGCAGAGAC for the nL mutation.

### RNA In situ hybridization (ISH) and Nanobody Immunohistochemistry in brain tissue

Embryonic brains were fixed in 4% PFA overnight and cut to 10 mm thickness as explained above. 10 mm Cryo-sections were pre-treated using the RNAscope Universal Pretreatment Kit (Advanced Cell Diagnostics) as indicated by the manufacturers. ISH were performed using the RNAscope Fluorescent Multiplex Reagent Kit as it was performed for cultured neurons (see above). The target gene (Ten4) was detected with the probe: Mm-Tenm4-C1 (RNAscope; Cat#555491). Following ISH, sections were immunostained O/N using the NanoTen4 conjugated to AlexaFluor633 (1:50) (generated and validated in this manuscript). NanoTen4 was diluted in 2% BSA, 0.3% Triton X-100, PBS. Nuclei were counterstained with DAPI, and sections were mounted with Mowiol. Images were acquired using a Zeiss LSM880 confocal laser scanning microscope using a 20x, objective and 2 Airy disk pinhole, and analysed with ImageJ ^110^ software.

### Model building and refinement

All model building was performed in Phenix ^113^ (version 1.20.1-4487) using the dock in map and real space refinement functions. Refinement in Phenix (using default constraints) and in COOT ^114^ was performed iteratively. Initial model generation was performed using PHYRE2^115^ inputting the Ten2 full ectodomain sequence (see vectors and cloning) for each splice site variant. Regions of this initial model not present in the cryo-EM density were trimmed after initial docking in ChimeraX ^116^. All figure panels showing structures or maps were made using ChimeraX ^116^. Cryo-EM density maps shown in Figure 1 and Extended Data Fig. 1, and also the density maps used for model building, were sharpened with deepEMhancer ^117^ (using the HighRes model). Model refinement statistics were reported with Phenix comprehensive validation using Molprobity ^118^ (see Table S2). The difference map between the Ten2 A1B1 and Ten2 A0B0 density maps was generated using the Surface colour volume tool in ChimeraX. The Ten2 A1B1 map was coloured by the volume data values of the Ten2 A0B0 map. The maximum value (0.25, depicted as blue) corresponds to voxels that present the same values in both maps. The minimum value (0, depicted as red) corresponds to voxels that are present in the Ten2 A1B1 map but not in the Ten2 A0B0 map. Note that 0.25 is the contour level used for all maps.

### Analysis of published single cell RNASeq datasets

Single cell RNAseq data for cortex samples (Fig. 4 and Extended Data Fig. 5) were obtained from the published NCBI Gene Expression Omnibus with accession number GSE153164^43^. For the single cell RNA-seq data for dissociated cortical neurons (Fig. 7 and Extended Data Fig. 7) we used a published NCBI Gene Expression Ommibus with accession number GSE271794 ^119^. We used the same UMAP coordinates and metadata information with the cluster categorization provided by the authors. We used the R package Seurat (v.4.0.1) to perform all the analysis.

### Nanobody discovery

We generated anti-Ten4 nanobodies by immunising one llama with the purified ternary complex of the mouse Teneurin4 ectodomain, the mouse Latrophilin1 Lec-Olf domains, and the mouse FLRT1 (mTen4-mFLRT1-mLphn/ADGRL1) as previously described ^120^. Briefly, a llama was immunized 6 times with purified ternary complex. Peripheral blood lymphocytes were collected after the final boost, RNA was purified, and cDNA created which served as template to generate a nanobody phage library in the pMESy4 vector encoding a His^6^-tag and EPEA-tag at the C-terminal end of the protein. After two rounds of phage display selection on the ternary complex or on mTen4 only, and screening by ELISA using established protocols ^120^. 13 different families, classified according to the sequences of the third complementarity determining region (CDR3), were discovered. These families were further screened using an optimised ELISA protocol (described above under Enzyme Linked Immunosorbent Assay (ELISA)) and one anti-Ten4 nanobody (CA20030) was selected as the best binder, which was used throughout this manuscript. The CDR loop sequences of CA20030 are GRTISNA (CDR1), SYSGRF (CDR2), and GSWSAIPNPSDYDYWGQG (CDR3). The structural model of Anti-Ten4 nanobody (CA20030) shown in Extended Data Fig. 4A,B was generated using Alphafold3 ^121^.

### Quantification and Statistical Analysis

Statistical analyses were performed using GraphPad Prism version 10, employing a one/two-tailed unpaired Student’s t test, and one-way ANOVA test with Tukey’s post hoc analysis when comparing multiple groups. For the analysis of the fraction of cells and the cumulative frequency distribution, we used chi-square contingency analysis and the Kolmogorov-Smirnov test, respectively. p values represent ∗p ≤ 0.05, ∗∗p ≤ 0.01, ∗∗∗p ≤ 0.001 and ∗∗∗∗p ≤ 0.0001. All statistical tests and p values for each panel are explained in the figure legends. All data are presented as the mean ± SEM, whisker plots, violin plots, bar plots or dot plots. All sample sizes and definitions are provided in the figure legends.

## Extended Data Figures

**Extended Data Fig 1.**
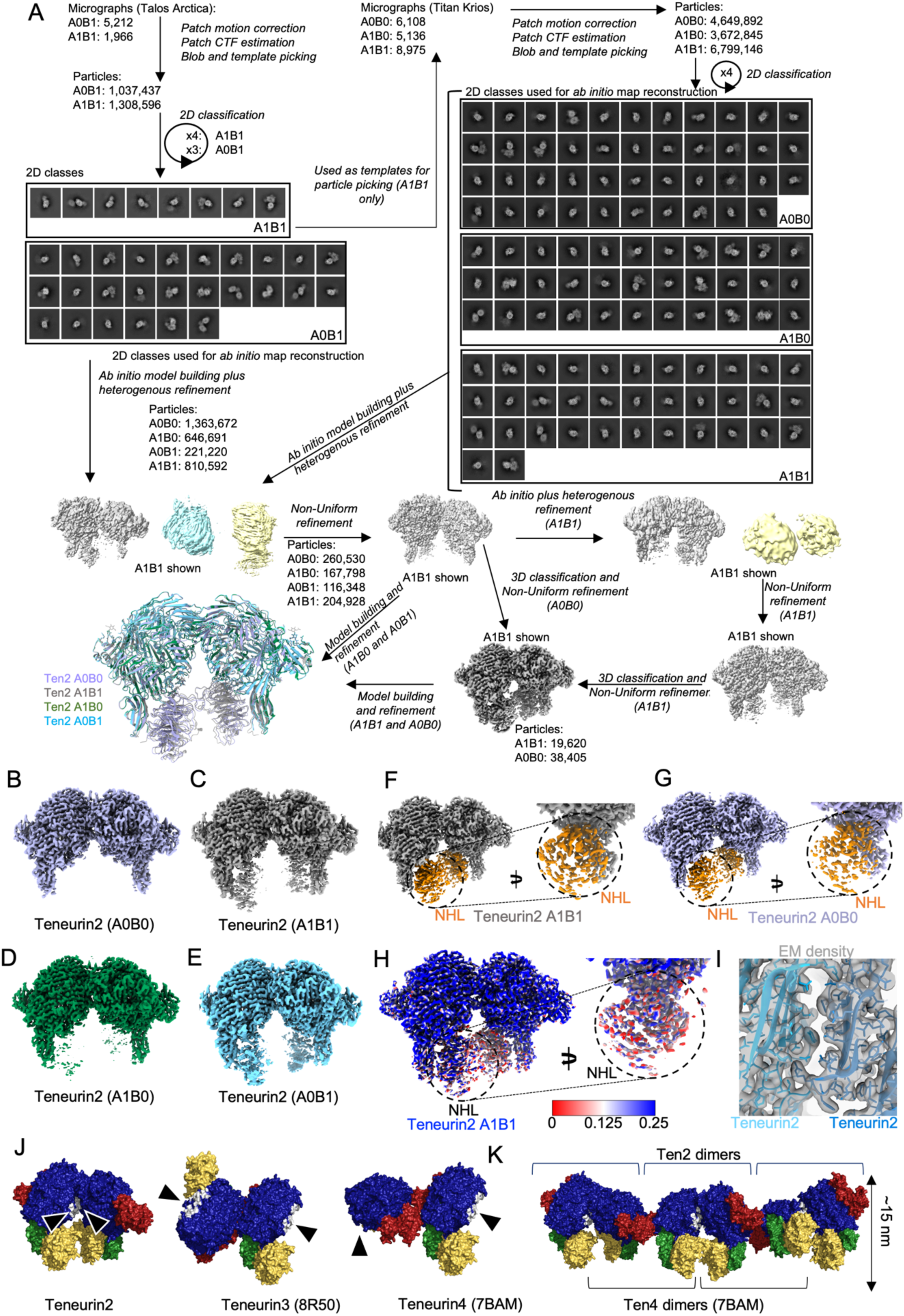
Single-particle Cryo-EM data analysis. **A**: Cryo-EM processing strategy and workflow in cryoSPARC. **B-E:** Resulting cryoEM maps for the four murine Ten2 homodimers. **F,G:** Resulting dimer maps for isoforms A1B1 (panel F) and A0B0 (panel G) of Ten2, derived using ∼15% of the particles after 3D classification and non-uniform refinement (see panel A). The NHL domain (corresponds to orange areas in the map) is better defined after this processing step. **H:** The map of the Ten2 A1B1 isoform is depicted (equivalent to that shown in panel F), but coloured according to how it differs from the map calculated for the A0B0 isoform (panel G). We used the surface colour volume tool and the colour by volume data values option in ChimeraX ^116^ to do this. The maximum value (0.25, depicted as blue), corresponds to voxels that present the same values in both maps. The minimum value (0, depicted as red), corresponds to voxels that are present in the Ten2 A1B1 map but not in the Ten2 A0B0 map. 0.25 is the contour level used for all maps. Most differences are found in the region corresponding to the NHL domain, which is less defined in the map for the A0B0 isoform. **I:** Zoomed-in view of the Teneurin2 dimer interface. The EM density map (grey) and model (shades of blue) are show. **J:** Comparison of the Ten2 dimer structure and previously published mammalian Teneurin dimers, which dimerise via different surfaces. Individual domains are coloured as follows: YD-shell: navy. NHL domain: yellow. FN-plug domain: green. C-terminal ABD and Tox-GHH domains: red. The Ten2-Ten2 interaction surface is coloured in white and indicated by arrow heads. The equivalent homologous surfaces on Ten3 and Ten4 are indicated in the relevant structures. **K:** Alternating the interfaces of the Ten2 and Ten4 dimer structures (PDB 7BAM ^9^) produce arrays. Ten2 structures are shown as surface views, coloured as in panel J.

**Extended Data Fig. 2:**
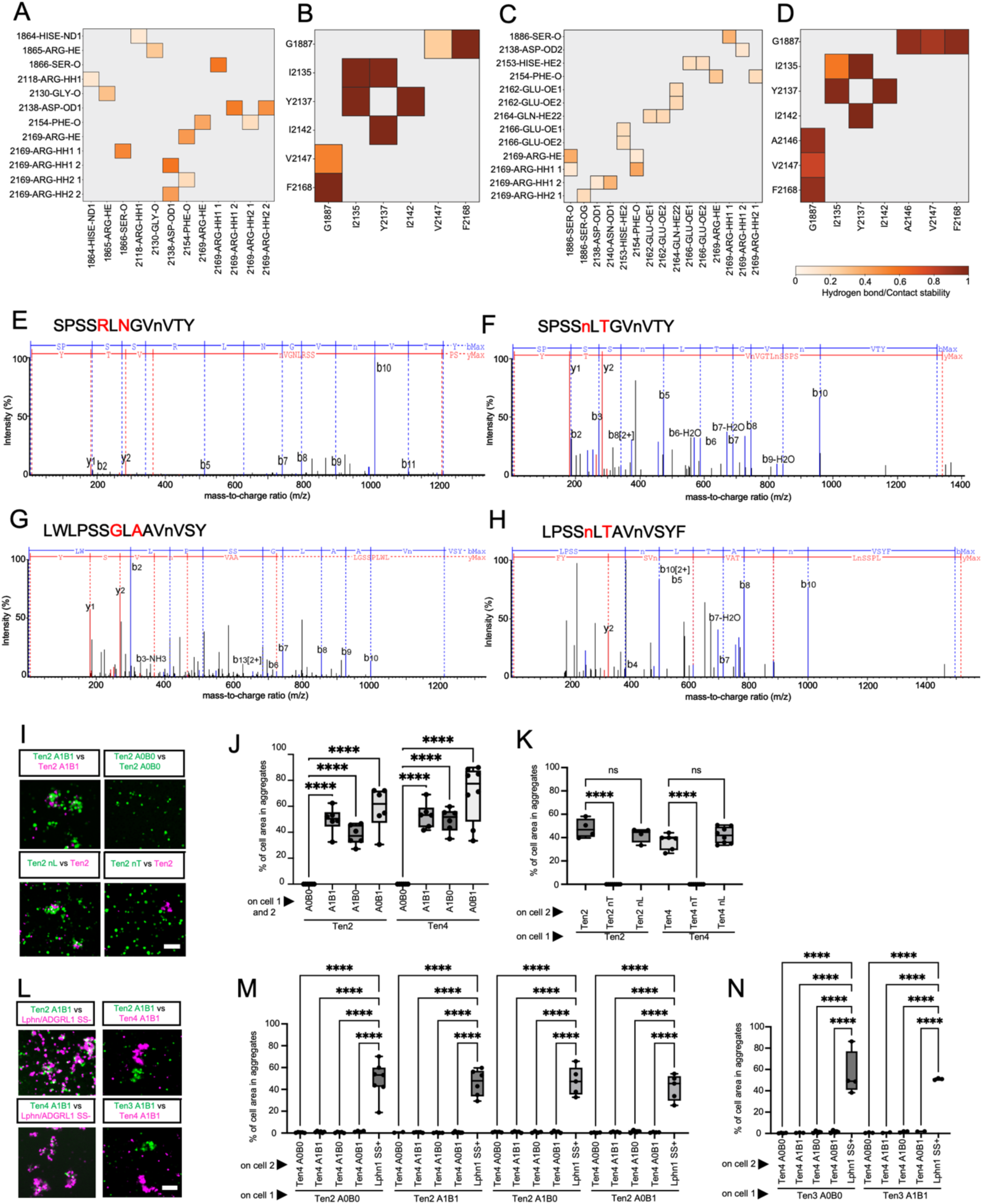
Ten2 interface analysis, mutant validation and cell aggregation. **A-D**: Summary of key residues that contribute to hydrophilic and hydrophobic interactions (red = most stable, white = least stable) in the Ten2-Ten2 interface, during MD simulation of the Ten2 dimer structure. Results are plotted after backbone constrained simulation (panels A, B) and after unconstrained simulation (panels C,D). **E-H:** We used liquid chromatography-mass spectrometry (LC-MS) to reveal N-linked glycosylation sites in chymotrypsin-derived Ten2 and Ten4 ectodomain peptides. Chymotrypsin cuts after F, W, Y. In the nT mutants we we introduced an N-linked glycan at positions G1887N (Ten2) and R1881N (Ten4) by mutating these sites to asparagine (N), followed by a non-proline residue, followed by a threonine (T). The mass spectrometry analysis confirms this. Results are shown for Ten2 WT (panel E), Ten2 nT (panel F), Ten4 WT (panel G) and Ten4 nT (panel H), focusing on the peptide that is mutated in the nT mutants. The mutated residues are highlighted in red. A lower-case n (instead of upper-case N) indicates that the site was indeed glycosylated. **I-K:** Cell-based aggregation results showing different levels of *’in trans’* binding for different Teneurin constructs. N>4 replicates. 6 pictures per replicate. **L-N:** Cell-based aggregation results showing no aggregation between the different Teneurin4 homologues Ten2, Ten3 and Ten4. N>2 replicates. 6 pictures per replicate. Representative images are shown in L. n.s. = not significant. ****p < 0.0001. One-way ANOVA test with Tukey’s post hoc analysis (J,K,M,N). Scale bars represent 100 μm (I,L).

**Extended Data Fig. 3.**
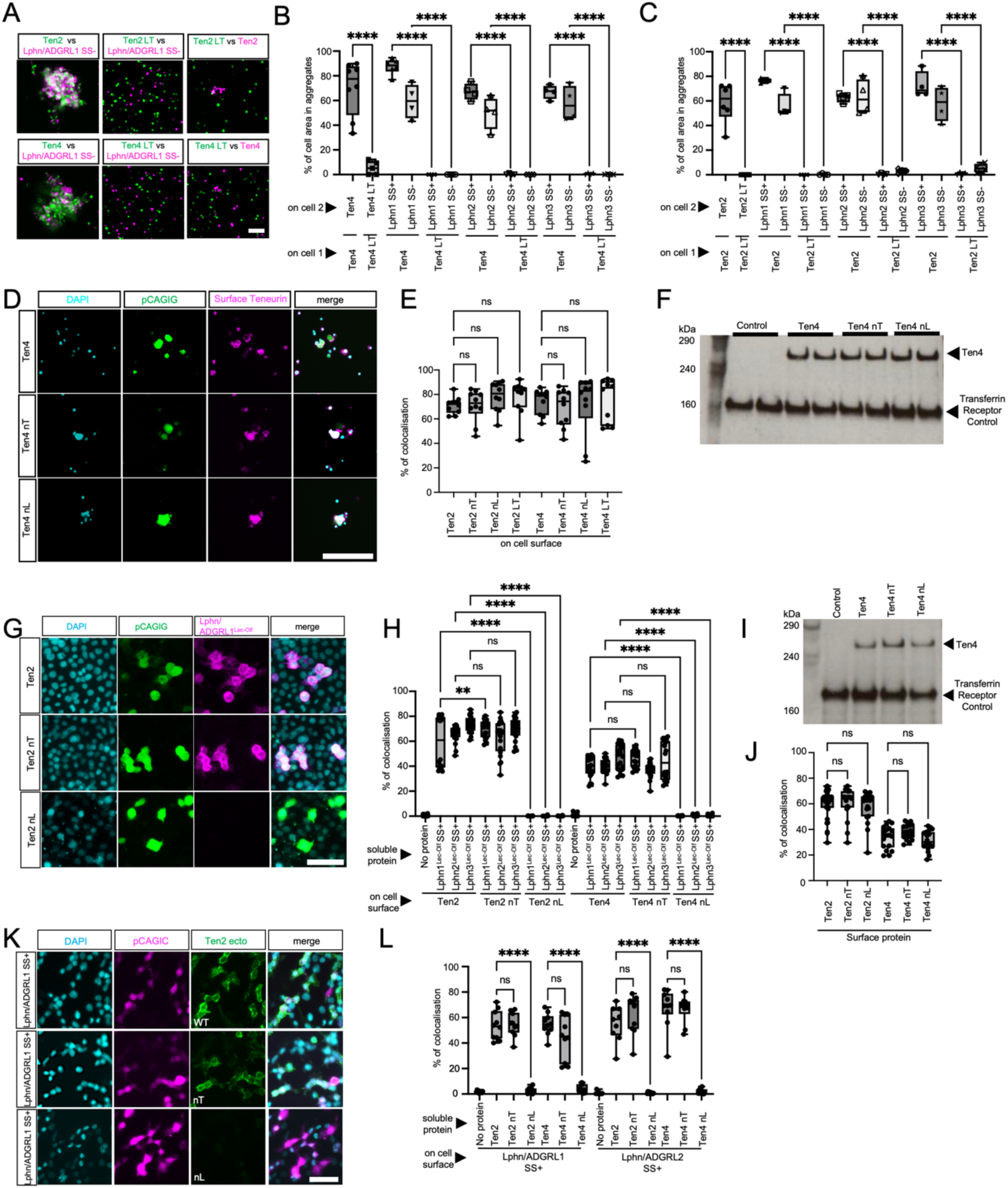
Cell aggregation and cell-based binding studies. **A-C:** Cell-cell aggregation results and representative images showing the Teneurin LT mutant ^3^ binding capabilities. N=4 replicates. 6 pictures per replicate. **D:** Representative images of a cell-surface staining experiment to assess protein expression levels at the cell surface of non-permeabilised K562 cells. **E:** Quantification of the surface expression levels of Ten2 and Ten4 WT, LT, nT and nL in K562 cells. Immunostaining against the extracellular HA-tag was used to assess surface presentation of these constructs. N=2 replicates. 10 pictures per replicate. **F:** Western blot results after the cell surface biotinylation and extraction of all membrane-bound proteins on K562 cells electroporated with either empty pCAGIG, Ten4 WT, Ten4 nT or Ten4 nL with an HA-antibody. The level of Transferrin Receptor in the samples was used as a loading control. **G,H:** Results of a cell-binding assay in which soluble Latrophilin Lec-Olf (magenta) is added to cells expressing Ten2/4 WT, nT or nL (green). Representative images are shown. N=3 replicates. 10 pictures per replicate. **I,J:** Assessment of the surface expression levels of Ten2 and Ten4 WT, nT, LT and nL in HEK293T cells, using analogous methods as panels D-F. N=2 replicates. 10 pictures per replicate. **K-L:** Results of cell-binding experiments as shown in panel G and H but using soluble Ten2 or Ten4 ectodomains (WT, nT or nL) with Latrophilin-expressing cells (magenta). N=1 replicate. 10 pictures per replicate. Representative images are shown. n.s. = not significant. ****p < 0.0001. One-way ANOVA test with Tukey’s post hoc analysis (B,C,E,H,J,L). Scale bars represent 100 μm (A,D,G,K).

**Extended Data Fig. 4.**
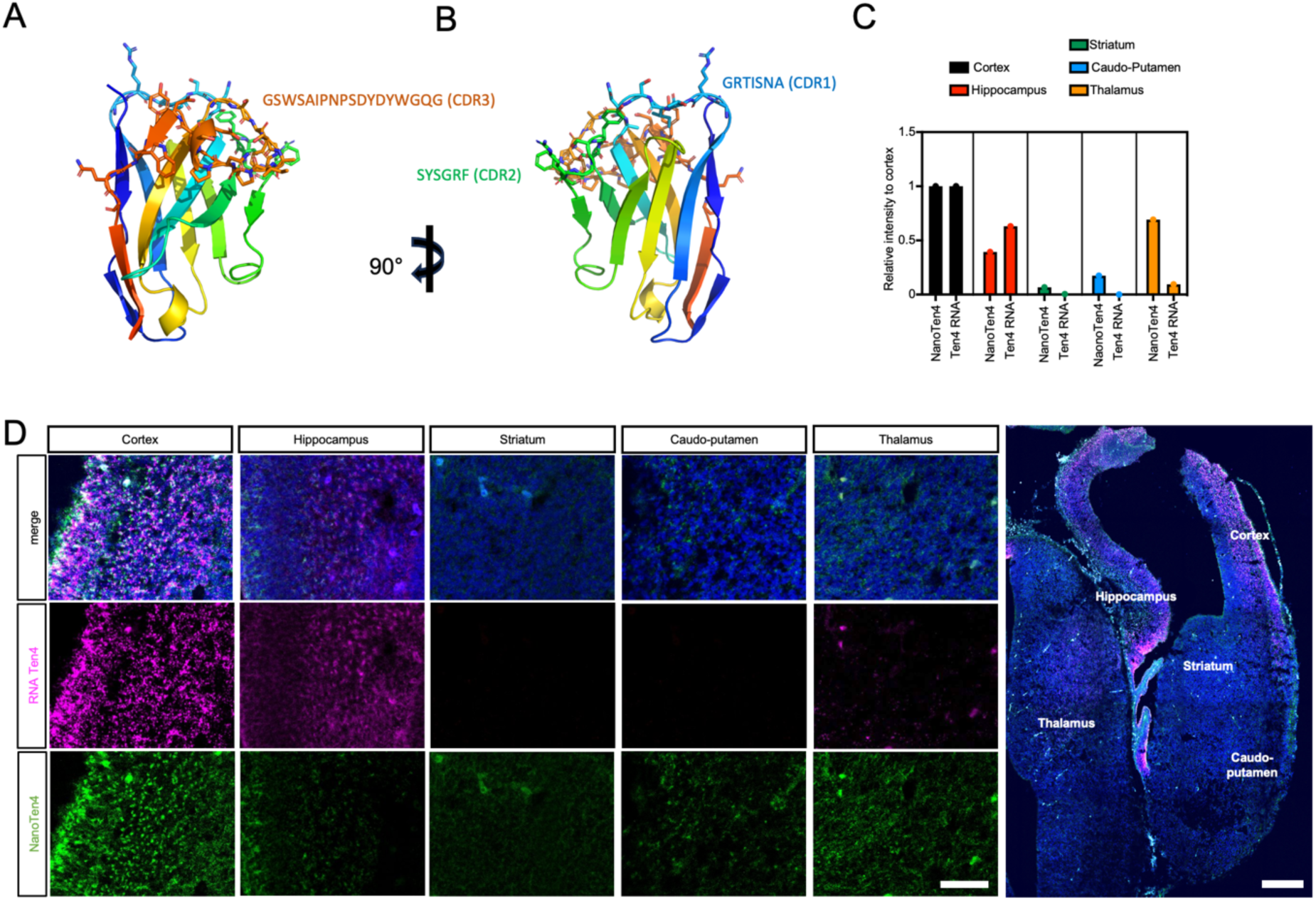
NanoTen4 and Ten4 *in situ* hybridisation analysis. **A,B**: Alphafold3^121^ structural model of anti-Ten4 nanobody (CA20030) with the CDR loop sequences highlighted and shown as sticks. The rest of the model is shown as a cartoon. **C**: Quantification of experiment shown in panel D. The NanoTen4 labelling and the presence of Ten4 mRNA largely correlate, suggesting that NanoTen4 labels Ten4 protein *in situ*. **D:** Different areas of an E15.5 murine brain slice, labelled with NanoTen4 and for Ten4 mRNA, were imaged and labelling area was quantified. Representative images are shown. N=3-4 sections/group from 2 mice. Scale bar represents 300 μm (A) and 50 μm inset (A).

**Extended Data Fig. 5.**
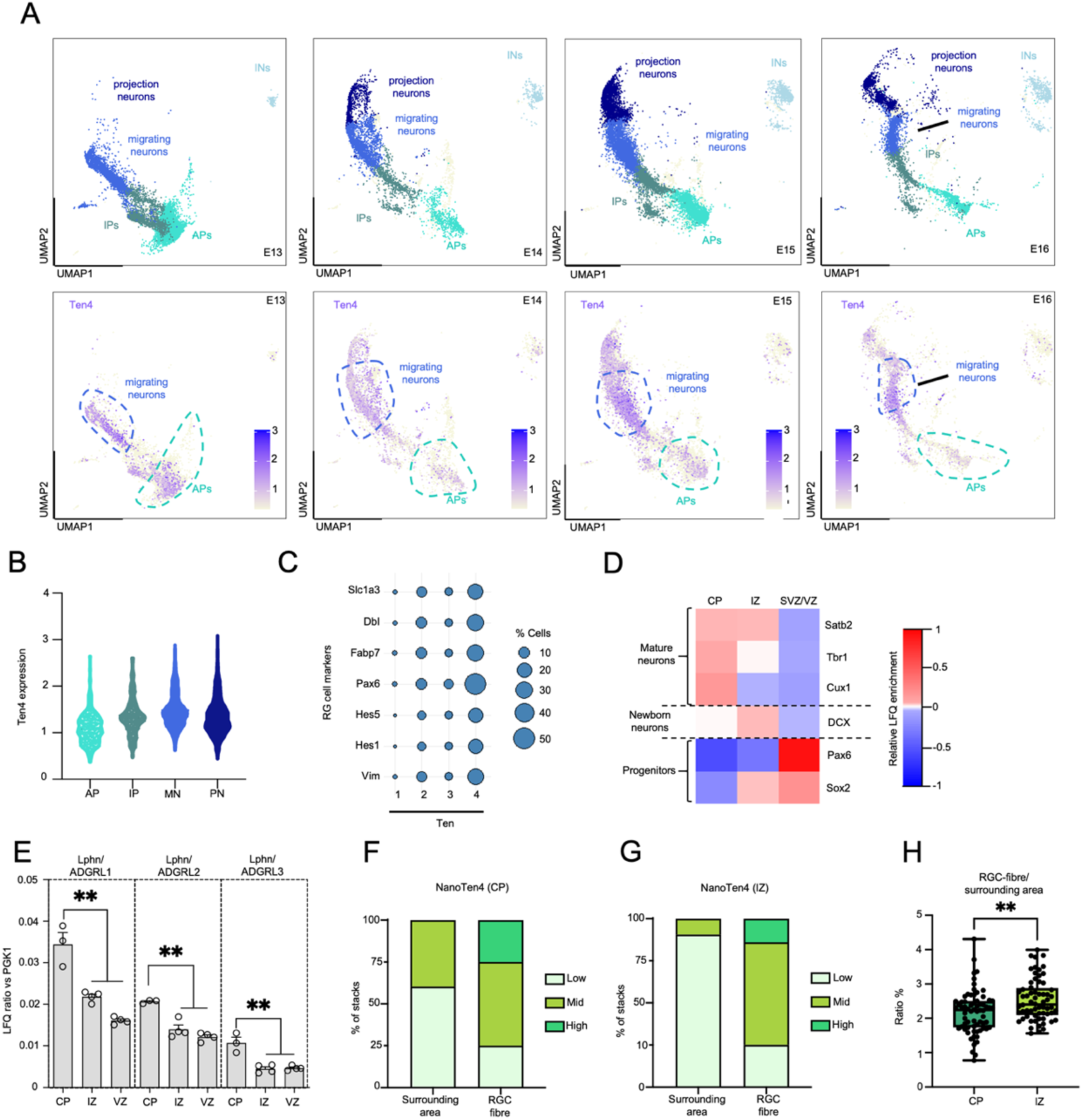
Ten4 expression analysis. **A**: UMAP visualizations of published single-cell data from E13, E14, E15, and E16 mouse cortex (GSE153164)^43^ and corresponding plots showing Ten4 mRNA expression. **B:** Volcano plot showing the distribution of Ten4-positive cells in apical progenitors (AP), intermediate progenitors (IP), migrating neurons (MN), and projection neurons (PN) at E15. **C:** Dot plot showing that the majority of radial glial cells (RGC) express Ten4. RGC cells are defined by expressing: Slc1a3, Dbl, Fabp7, Pax6, Hes5, Hes1 and Vim. **D:** Analysis of neuronal and progenitor markers in mass spectrometry data confirms good separation of the cortical layers indicated. CP=cortical plate, IZ=intermediate zone, SVZ/VZ=subventricular/ventricular zones. **E:** Analysis of the expression of Latrophilins (Lphns/ADGRLs) in different cortical layers using mass spectrometry. N=3-4 samples per group. **F,G:** We determined thresholds for low, mid, and high NanoTen4 labelling intensity using the 25 and 75 percentiles for the values of NanoTen4 staining in the areas within a 1.5 μm radius of the BLBP-positive fibres in the CP as thresholds. We plotted the data separately for areas within a 1.5 μm radius of the BLBP-positive fibres, and outside of these areas (’surrounding area’) for both CP and IZ. **H:** Comparison of the ratio between NanoTen4 staining along radial glia cell fibres versus the surrounding area, comparing quantifications for the cortical plate versus the intermediate zone. **p < 0.01. One-way ANOVA test with Tukey’s post hoc analysis (E). Student’s t test (H).

**Extended Data Fig. 6.**
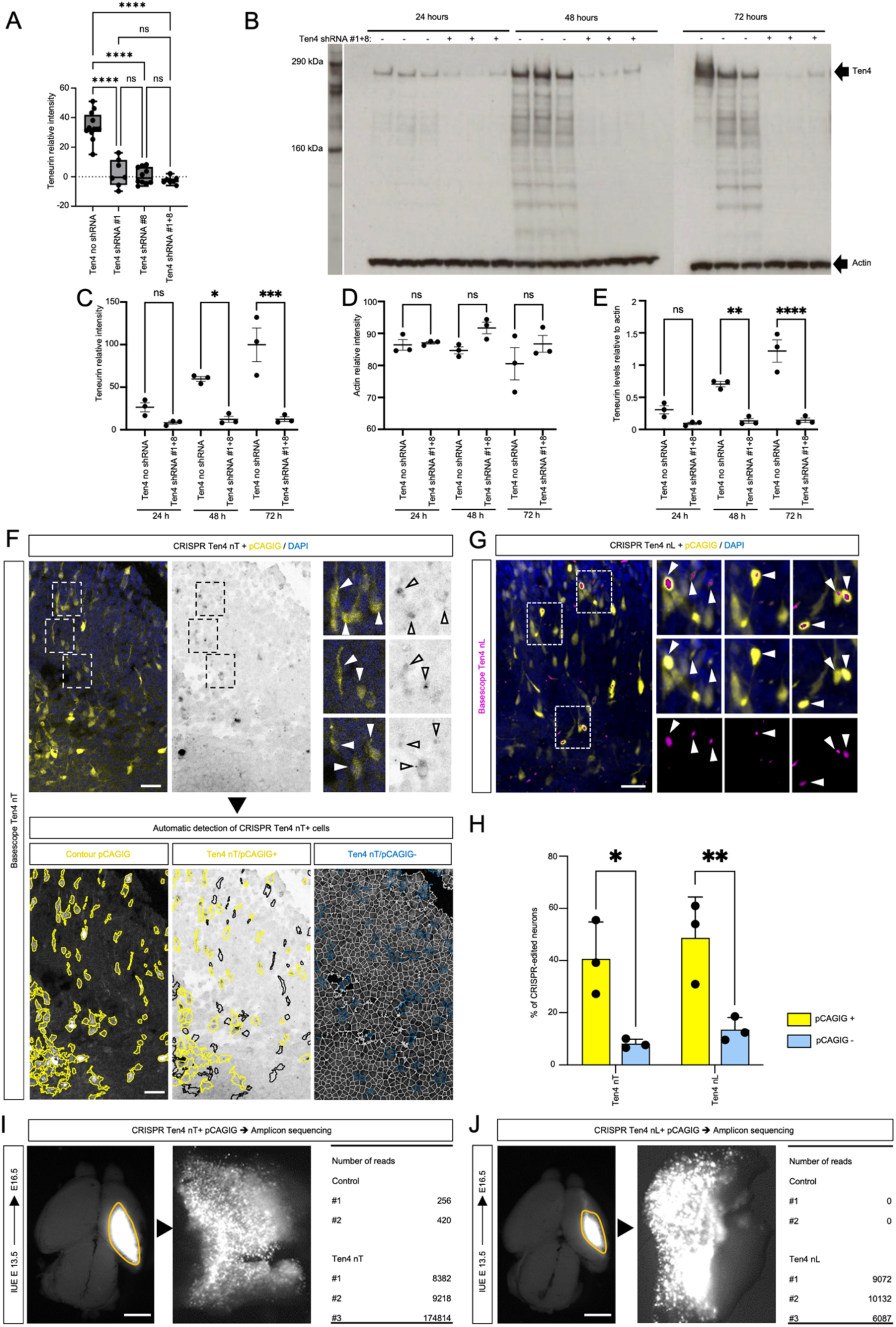
Validation of reagents used for *in vivo* analysis. **A**: We tested the efficacy of shRNA constructs designed to knock down Ten4 expression with Ten4-overexpressing HEK293T cells. In these experiments, HA-tagged Ten4 is co-transfected with plasmids encoding two shRNA constructs, either separately or alone (#1 and #8). Ten4 expression levels were quantified at 72h post transfection, using anti-HA western blotting. **B:** Based on results in panel A, we focused on using both shRNA constructs (#1 and #8), and tested their efficacy at different time points (24 hours, 48 hours, 72 hours). Actin was used as a loading control. The knock-down effect was present also after 72 hours, which is the length of an IUE experiment *in vivo*. **C-D:** Quantification of the intensity of protein bands (Ten4 or actin) relative to background in the Western blot shown in panel B. Box and whiskers plot. N=3 replicates. **E:** Quantification after normalisation, combining data in panels C+D. **F-G:** Basescope^TM^ labelling and automated analysis validating the successful mutation of endogenous Ten4 genes using CRISPR/Cas9 in pCAGIG (GFP) expressing cells. **H:** Quantification from panels F-G. N=3 embryos. **I,J:** Amplicon sequencing confirms the presence of CRISPR/Cas9-edited cells expressing mutant Ten4 mRNA. *p < 0.05. **p < 0.01. ***p < 0.001. ****p < 0.0001. One-way ANOVA test with Tukey’s post hoc analysis (C-E). Student’s t test (H). Scale bar represents 900 μm (I,J).

**Extended Data Fig. 7.**
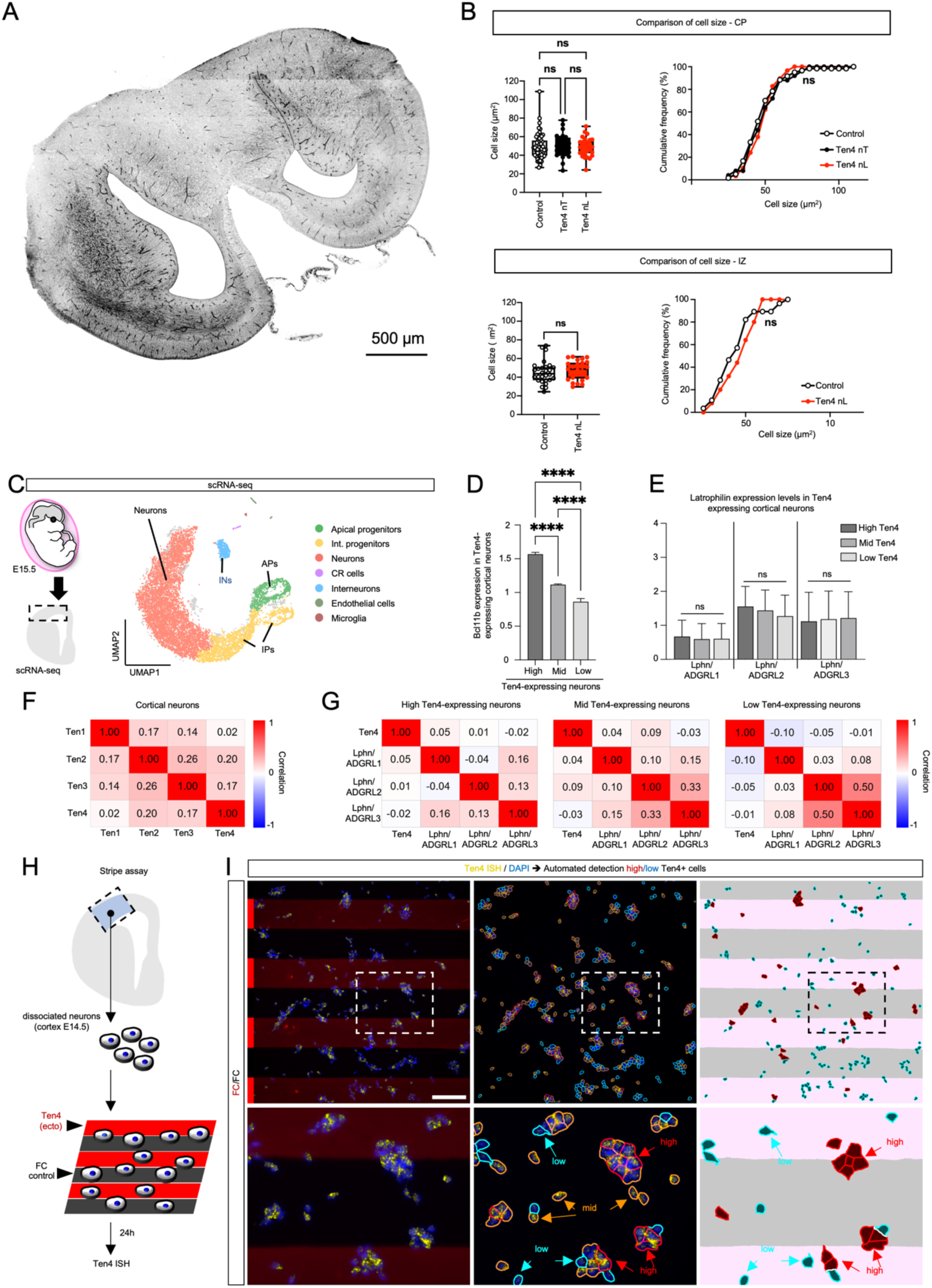
Shadow labelling suggests no change in cell size for Ten4 mutant neurons, and stripe assay method using cortical neurons. **A**: Overview of a whole fixed and shadow-labelled brain slice. **B:** Whisker plot and cumulative frequency distribution plot for the quantification of the cell sizes of migrating neurons (mutant and control) in the cortical plate (CP) and intermediate zone (IZ). CRISPR/Cas-9 edited cells containing Ten4 mutations (nT and nL) and WT control cells were analyzed. Note that there are almost no nT mutants left in the IZ at the end of the experiment and therefore only the nL mutant was analyzed. **C:** Schematic and UMAP of an scRNA-seq dataset for murine cortex at E15.5. Cell types are coloured as indicated in the legend (data source: GSE271794^119^) **D:** Bcl11b expression in cortical neurons grouped according to their Ten4 expression levels, as defined in Fig. 7E. **E:** Latrophilin mRNA expression levels in these different Ten4-expressing populations. **F:** Heatmap showing correlations for the mRNA expression levels of different Teneurin isoforms in cortical neurons (same data source as panel C). Red = positive high correlation, white = no correlation, blue = negative high correlation. **G:** As panel E but showing correlations for Ten4 and Latrophilin mRNA expression. We analysed these correlations separately for cells expressing different Ten4 levels. **H:** Overview of the stripe assay method. **I:** Snapshots taken from the automated analysis pipeline for Ten4 stripe assays. We used CellProfiler^111^ to outline cells according to the Ten4 mRNA transcript levels detected using RNAscope (red = high, orange = mid, cyan = low). The percentage of high and low Ten4-expressing cells located on red stripes (highlighted in pink after thresholding) was quantified in ImageJ using a custom macro from a previous study^3^. As an example, Fc/Fc control data are shown. n.s. = not significant. * p<0.05. One-way ANOVA test with Tukey’s post hoc analysis (Cell size, panel B,D,E) and Kolmogorov-Smirnov test (cumulative frequency distribution, panel B). Scale bars represent 500 μm (A) and 100 μm (I).

Supplementary table 1 related to Figure 1 and Extended Data Fig. 1. Cryo-EM processing statistics for the 3D maps.

|  | Teneurin2 A0B0<br>EMDB: EMD-51022 | Teneurin2 A1B1<br>EMDB: EMD-50975 | Teneurin2 A1B0<br>EMDB: EMD-50976 | Teneurin2<br>A0B1<br>EMDB: EMD-51021 |
| --- | --- | --- | --- | --- |
| Magnification | 105,000 | 105,000 | 105,000 | 150,000 |
| Voltage (kV) | 300 | 300 | 300 | 200 |
| Dose per frame (e<sup>-</sup>/Å<sup>2</sup>) | 1.04 | 1.06 | 1.04 | 1 |
| Electron exposure (e<sup>-</sup>/Å<sup>2</sup>) | 41.503 | 42.303 | 41.503 | 40 |
| Defocus range (µm) | -1 to -2 | -1 to -2 | -1 to -2 | -1.5 to -2.75 |
| Pixel size (Å) | 0.83 | 0.83 | 0.83 | 0.94 |
| Symmetry imposed | C2 | C2 | C2 | C2 |
| Number of micrographs | 6,108 | 8,975 | 5,136 | 5,212 |
| Initial number of particles | 4,649,892 | 6,799,146 | 3,672,845 | 1,037,437 |
| Final number of particles | 38,045 | 19,620 | 167,798 | 116,348 |
| Map resolution (FSC 0.143) | 2.82 Å | 2.80 Å | 2.55 Å | 3.48 Å |

As reported by cryoSPARC

Supplementary table 2 related to Figure 1 and Extended Data Fig. 1. Refinement statistics for the atomic models.

|  | Teneurin2<br>A0B0<br>PDB: 9G42 | Teneurin2<br>A1B1<br>PDB: 9G2F | Teneurin2<br>A1B0<br>PDB: 9G2H | Teneurin2<br>A0B1<br>PDB: 9G41 |
| --- | --- | --- | --- | --- |
| Chains | 4 | 4 | 2 | 2 |
| Total number of atoms | 27,564 | 28,160 | 22,241 | 22,275 |
| Number of residues | 3,444 | 3,504 | 2,732 | 2,740 |
| R.m.s deviation bond lengths (Å) | 0.004 (0) | 0.004 (0) | 0.003 (0) | 0.003 (0) |
| R.m.s deviation bond angles (°) | 0.713 (10) | 0.701 (5) | 0.595 (0) | 0.594 (0) |
| Ramachandran plot favoured (%) | 96.88% | 96.46% | 97.39% | 97.14% |
| Ramachandran plot allowed (%) | 3.12% | 3.54% | 2.61% | 2.86% |
| Ramachandran plot outlier (%) | 0% | 0% | 0% | 0% |
| Rotamer outliers (%) | 0.90% | 1.45% | 0.79% | 0.67% |
| Clash score | 3.83 | 3.68 | 3.69 | 3.52 |
| Molprobity score | 1.36 | 1.51 | 1.28 | 1.30 |
| CC model vs data (mask) | 0.84 | 0.85 | 0.83 | 0.78 |

As reported by PHENIX comprehensive cryo-EM validation and Molprobity.

